# The Initiation Knot is a Signaling Center Required for Molar Tooth Development

**DOI:** 10.1101/2020.04.09.033589

**Authors:** Isabel Mogollón, Jacqueline E. Moustakas-Verho, Minna Niittykoski, Laura Ahtiainen

**Affiliations:** Cell and Tissue Dynamics Research Program, Institute of Biotechnology, University of Helsinki Finland; Organismal & Evolutionary Biology Research Program, University of Helsinki Finland

**Author notes:** Corresponding author L.A., Cell and Tissue Dynamics Research Program, Institute of Biotechnology/Helsinki Institute of Life Science, P.O. Box 56 (Viikinkaari 5), FIN-00014 University of Helsinki, Finland.

**Keywords:** cell division, migration, embryonic development, tooth, signaling center, Wnt, Shh

## Abstract

Signaling centers, or organizers, regulate many aspects of embryonic morphogenesis. In the mammalian molar tooth, reiterative signaling in specialized centers called enamel knots (EKs) determine tooth patterning. Preceding the first, primary EK, a transient epithelial thickening appears whose significance remains debated. Here, using tissue confocal fluorescence imaging with laser ablation experiments, we show that this transient thickening is an earlier signaling center, the molar initiation knot (IK) that is required for the progression of tooth development. IK cell dynamics manifest the hallmarks of a signaling center; cell cycle exit, condensation, and eventual silencing through apoptosis. IK initiation and maturation are defined by the juxtaposition of high Wnt activity cells to *Shh*-expressing non-proliferating cells, the combination of which drives the growth of the tooth bud, leading to the formation of the primary EK as an independent cell cluster. Overall, the whole development of the tooth, from initiation to patterning, is driven by the iterative use of signaling centers.

## Introduction

In recent years, advances in 3D and live tissue imaging have brought new understanding of the cell level behaviors that contribute to the highly dynamic stages of morphogenesis in ectodermal organs, such as hair and teeth (Kim, et al., 2017; Ahtiainen, et al., 2016; Ahtiainen, et al., 2014; Devenport and Fuchs, 2008). Despite shared morphological characteristics and conserved signaling (Biggs and Mikkola, 2014; Jernvall and Thesleff, 2000), it is becoming evident that signaling cues are interpreted into diverse cellular behaviors depending on the context, thereby defining different organ shapes and sizes already at early stages of organogenesis. Morphogenesis in ectodermal organs is regulated by epithelial signaling centers that form sequentially in specific spatiotemporal patterns and govern cell behaviors via secreted factors including hedgehog (Hh), Wnt, fibroblast growth factor (Fgf) and bone morphogenic protein (Bmp) family members (Jernvall and Thesleff, 2000; Dassule and McMahon, 1998).

Teeth have long served as a model organ to study mechanisms of embryonic development in tissue interactions and genetic regulation (Jernvall and Thesleff, 2000). Mice have two tooth types: large ever-growing incisors and multicuspid molars. Organogenesis in teeth is initiated at embryonic day (E)11 with an epithelial thickening called the dental lamina. It resolves into separate domains for incisor and first molar primordia, with a toothless diastema in between. The instructive potential resides initially in the epithelium and shifts to the mesenchyme at the bud stage. Epithelial budding starts at E12.5 followed by mesenchymal condensation leading to a mature bud at E13.5 (Jernvall and Thesleff, 2000). The molar primary enamel knot (pEK) signaling center appears at E13.5 in the late bud stage epithelium and matures into the enamel organ in the cap stage at E14.5 (Tummers and Thesleff, 2009). The pEK is silenced by apoptosis and sequentially followed by pairs of secondary enamel knots (sEK) that regulate cusp patterning (Jernvall, et al., 1994). Fate mapping studies have shown that the pEK clonally contributes to the buccal sEK, but may not contribute to the lingual counterpart (Du, et al., 2017).

The cellular events in early molar morphogenesis have remained largely uncharted, as emphasis has been on the bud stage and beyond. Recently, we identified a novel epithelial signaling center in the early developing incisor, called the initiation knot (IK), which drives local cell proliferation for epithelial budding (Ahtiainen, et al., 2016). The incisor IK shares transcriptional signatures with the incisor enamel knot (EK), which forms without clonal contribution from the IK (Du, et al., 2017; Ahtiainen, et al., 2016; Li, et al., 2016a). While the molar placode and EKs are known to share molecular markers, a similar signaling center in the molar as in the incisor has not been reported. However, previous studies using expression and histological analyses of molar morphogenesis prior to budding, have interpreted a transient epithelial thickening in the diastema anterior to the first developing molar as evidence for the presence of vestigial premolar teeth lost during murine evolution (Prochazka, et al., 2010).

To resolve the early events of molar morphogenesis, we use confocal fluorescence whole-mount live tissue imaging to elucidate the cellular and molecular dynamics of signaling centers and how they shape the tooth bud. We show that an IK signaling center is established in the molar placode and it remains an integral functional part of the developing bud. The molar IK arises by the juxtaposition of cells with high canonical Wnt activity to *Shh*-expressing G_1_/G_0_-phase cells. Molar early growth is dependent on the IK signaling center and interference of the function of this signaling center either mechanically, by laser ablation, or with specific modulators of relevant signaling pathways, abrogates bud proliferative growth and progression of tooth development. The IK positions the tooth in the growing mandible and is silenced by apoptosis as the pEK arises independently to drive further growth of the bud. The cellular and molecular dynamics of the IK signaling center control tooth development earlier than previously thought.

## Results

### A molar initiation knot is established in the placode and early bud in G_1_/G_0_ cells expressing signaling center markers

Cell cycle exit is an early hallmark of ectodermal placodes (Ahtiainen, et al., 2016; Ahtiainen, et al., 2014). The Fucci fluorescent cell cycle reporter system allows direct real-time follow-up of the progress of the cell cycle in individual cells in the developing tissue: When the cell is in G_1_/G_0_ phase the nucleus emits red fluorescence and upon transition to S/G_2_/M proliferative phase, the cell nucleus emits green fluorescence. We used confocal fluorescence microscopy of whole-mount mandibles of the Fucci cell cycle indicator transgenic mouse to characterize G_1_/G_0_ cell distribution in the developing molar. Transgenic Shh^GFP/+^ (Harfe, et al., 2004) and Fgf20^βGal/+^ expression were used to identify signaling centers from E11.5-E13.5 and EpCam immunofluorescence staining to visualize the epithelium.

At E11.5, G_1_/G_0_ phase cells were distributed throughout the dental lamina (Fig.1A). By E12.5 the G_1_/G_0_ cells were located mesially in the mature placode/early bud. At 13.0, the G_1_/G_0_ focus remained in the mesio-lingual part of the bud, close to epithelial surface and a new focus of G_1_/G_0_ cells appeared distally deep in the invaginating bud, in the presumptive pEK area. By E13.5, the early G_1_/G_0_ focus was lost with only a few cells remaining (Fig.1A). Quantification of G_1_/G_0_ cells in different stages showed a decrease in the G_1_/G_0_ cell number in the early focus from E12.5 to E13.5 (Fig.S1A). In parallel, G_1_/G_0_ cells corresponding to the pEK area emerged. For further functional analyses we verified that the budding morphogenesis and G_1_/G_0_ cell distribution were similar in *ex vivo* cultured whole mount explants to *in vivo* (Fig.S1B). Numbers of G_1_/G_0_ cells were also similar *in vivo* and in cultured tissue (Fig.S2C).

**Figure 1.**
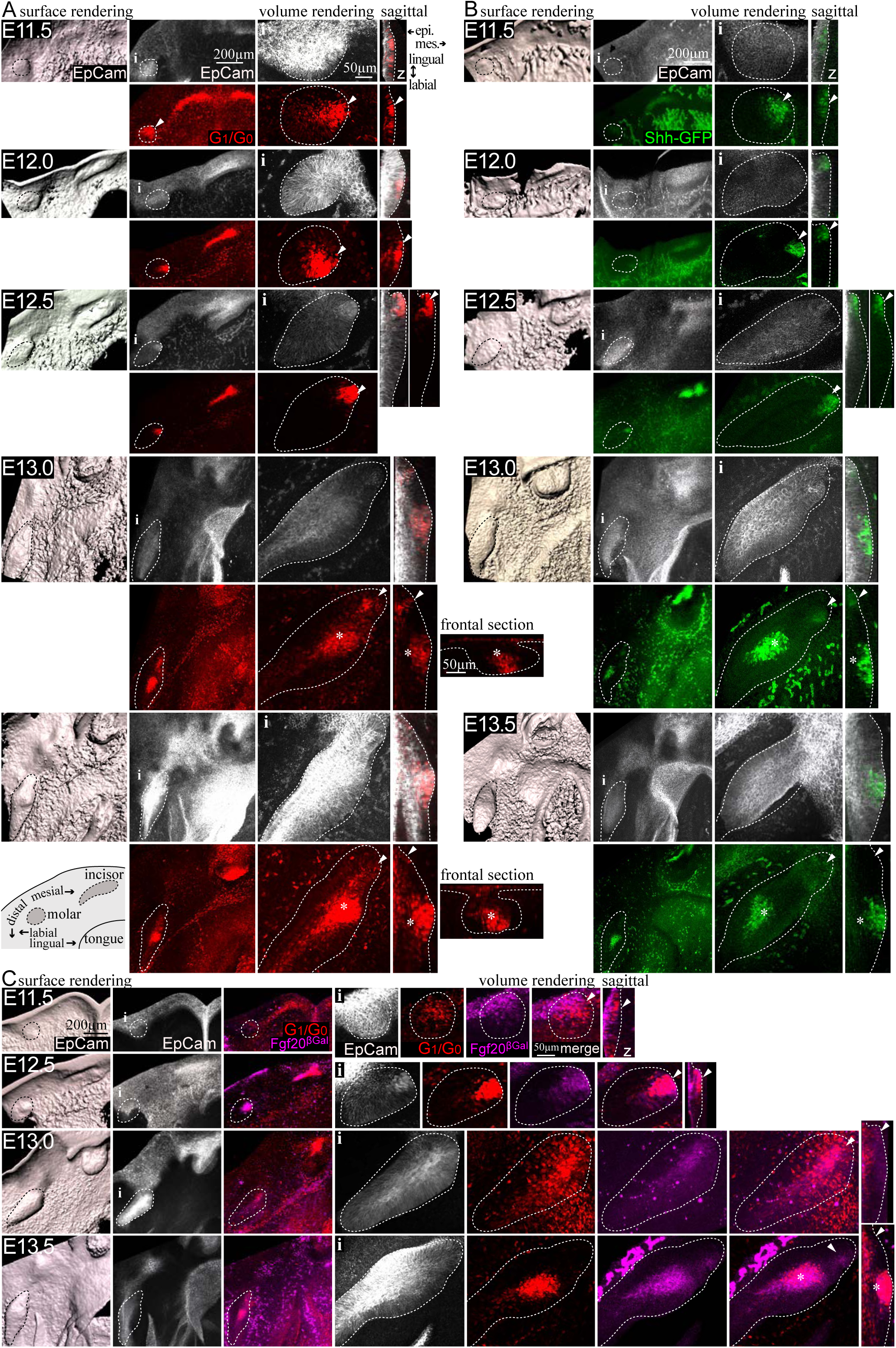
A molar initiation knot is established in the molar placode and early bud in G_1_/G_0_ cells positive for signaling center markers. Confocal fluorescence images of mouse embryonic mandibles of the cell cycle indicator (Fucci) for G_1_/G_0_ phase (red) and signaling center marker Shh^GFP^(green) and Fgf20^βGal^(βGal staining, magenta) Immunofluorescence staining of the epithelium (EpCam, grey, dotted line), early G_1_/G_0_ focus (arrowhead), presumptive primary enamel knot (pEK)(asterisk). Planar view from the mesenchyme toward the epithelium. (A) G_1_/G_0_ phase cells were present throughout the dental lamina and the molar placode, as incisor and molar resolved into separate domains at E11.5. At E12.5, G_1_/G_0_ cells formed a focus mesially in the molar early bud. This focus remained close to epithelial surface mesio-lingually. At E13.0, G_1_/G_0_ cells corresponding to the presumptive pEK emerged in the tip of the bud and condensed by E13.5. (B) Shh^GFP^ reporter and (C) Fgf20^βGal^ signaling center marker showed expression corresponding to G_1_/G_0_ foci throughout placode and bud morphogenesis and in the emerging pEK.

The Shh^GFP^ and Fucci G_1_/G_0_ reporters could not be combined as this often resulted in abnormal development of the craniofacial structures. However, the Shh^GFP^ reporter showed expression in the same areas as the G_1_/G_0_ foci throughout morphogenesis (Fig.1B): GFP+ cells were present throughout the placode at E11.5. At E12.5, they were located at the mesio-lingual side of the early bud close to the epithelial surface. By E13.5, GFP signal had disappeared almost completely in the early G_1_/G_0_ focus and appeared in the presumptive pEK area. DIG *in situ* hybridization with a probe specific for *Shh* in Fucci G_1_/G_0_ reporter mandibles showed exact colocalization of *Shh* with the Fucci reporter (Fig.S1E). The numbers of Shh^GFP^+ cells and Fucci G_1_/G_0_ showed similar distribution in the developing molar (Fig.S1A,D). Immunofluorescence staining for β-galactosidase (βGal), marking signaling centers in Fgf20^βGal/+^;Fucci G_1_/G_0_ embryos, showed colocalization of the markers from E11.5 in the early G_1_/G_0_ focus and through early bud stage (E12.5-E13.0) and in the pEK at E13.5 (Fig.1C).

Together these data confirmed the identity of the initial molar placode G_1_/G_0_ focus and corresponding focus in the mesio-lingual part of the developing molar bud epithelium as a signaling center. This early signaling center appeared prior to the pEK and thus we call this signaling center a molar initiation knot (IK).

### The molar IK is a functional signaling center driving the molar bud proliferative growth

Next, we studied cell proliferation in the developing molar. In the incisor, budding occurs via cell proliferation regulated by non-proliferative signaling centers (Ahtiainen, et al., 2016), whereas cell rearrangements and migration together with Shh driven proliferation have been proposed as mechanisms for molar bud invagination (Li, et al., 2016b; Prochazka, et al., 2015; Dassule and McMahon, 1998). To dissect the IK contribution to the molar bud, we first studied cell proliferation with Fucci S/G_2_/M and G_1_/G_0_ cell cycle indicators in fixed whole-mount mandibles. We then imaged whole-mount mandible explant cultures using live tissue confocal microscopy, which allowed us to follow the developing bud in a single-cell resolution.

Live tissue confocal microscopy of the Fucci G_1_/G_0_ reporter, for visualization of the molar IK and pEK cells, and K17-GFP reporter to follow the shape of the epithelial bud from E12.5+16h, confirmed that the IK cells stay an integral part of the developing bud (Fig.2A, Fig.S2A). Observing proliferation patterns with the Fucci reporters showed that during early initiation, at E11.5, G_1_/G_0_ cells comprised the placode and S/G_2_/M cells were evenly distributed throughout the oral epithelium (Fig.2B,C). By E12.5, S/G_2_/M cells appeared posterior to the IK in the maturing placode. From E12.5 to E12.75, there was a sharp increase in S/G_2_/M cells throughout the bud epithelium, in both basal and suprabasal populations (Fig.2B,C). At E13.5, S/G_2_/M cells were present in the bud and surrounding the pEK area. Few IK G_1_/G_0_ cells still remained. Quantification of cell number showed very few proliferative cells in the placode at E11.5 (Fig.S2B). At E12.5, during the initiation of budding, there was a threefold increase in S/G_2_/M cell number and further a twofold increase at E13.0 with similar cell numbers in *ex vivo* cultures (Fig.S2B). Proliferation was concurrent with bud elongation and invagination (Fig.S2C).

**Figure 2.**
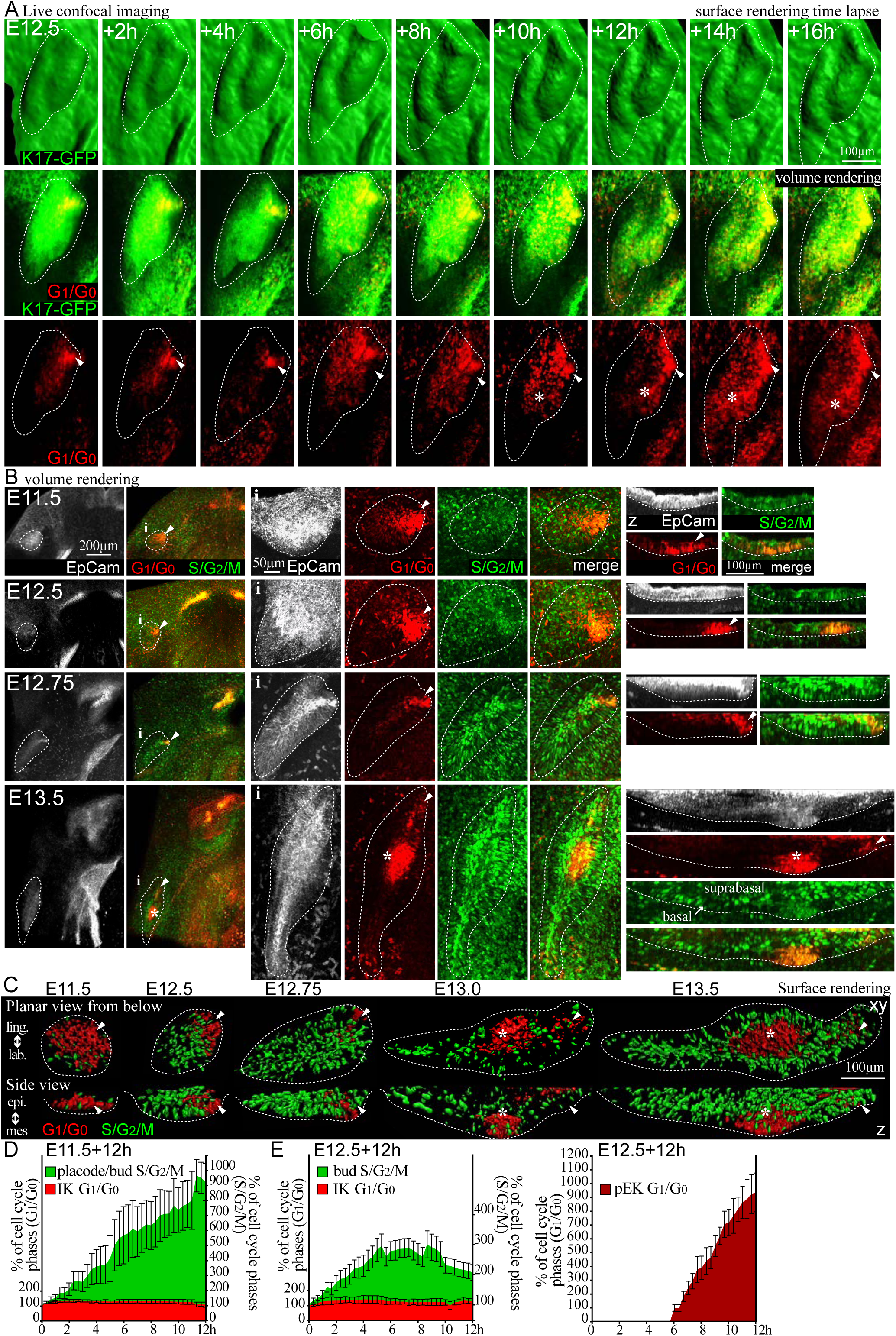
The molar IK is a functional signaling center driving budding via proliferation. (A) Still images of live tissue confocal microscopy of the Fucci G_1_/G_0_ reporter to visualize the molar IK cells and K17-GFP (green) reporter to visualize the borders and shape of the epithelial bud from E12.5+16h showed that the IK stays an integral part of the developing bud. (B) Confocal fluorescence images of whole mount explants Fucci G_1_/G_0_ (red), S/G_2_/M (green), epithelium (EpCam, white, dotted line), IK (arrowhead) and pEK (asterisk). Initially S/G_2_/M cells were seen throughout the oral epithelium and by E12.5, in the early bud posterior to the IK in both basal and suprabasal populations. IK and pEK cells remained in G_1_/G_0_ phase. (C) Surface rendering of the nuclei in G_1_/G_0_ and S/G_2_/M cell cycle phases in the developing molar placode/bud epithelium. (D) Quantification of cell cycle phases in live imaging E11.5+12h and (E) E12.5+12h molars (N=5, N=3, mean±SEM).

We next analyzed the contribution of individual cells in each cell population to the growing bud. Quantification of cell cycle phases with live imaging from E11.5+12h molars showed constant G_1_/G_0_ cell number in the placode (Fig.2D). A burst of cell proliferation from 4h onwards was seen in the emerging bud. This was specific to the tooth bud, as the contributions of G_1_/G_0_ and S/G_2_/M cells remained constant in the oral epithelium (Fig.2D, Fig.S2D). When we followed individual IK cells through the cell cycle from E11.5+12h, we observed some new G_1_/G_0_ cells appearing in the IK while two cells showed nuclear fragmentation (Fig.S2E, Mov.S1). None of the followed G_1_/G_0_ cells in the IK re-entered the cell cycle. Live imaging from the early bud stage onwards E12.0+12h, showed that more bud cells entered S/G_2_/M (Fig.S2F, Mov.S2). There was a respective increase in cell divisions throughout the bud as the bud grew. When we followed individual proliferating cells, of the 126 original S/G_2_/M cells followed, 25% went through cytokinesis, and divisions were observed throughout the bud (Fig.S2F, Mov.S2). Some IK cells showed nuclear fragmentation and were lost, while remaining IK cells stayed in G_1_/G_0_. Quantification of cell cycle phases from E12.5+12h showed that number of molar IK G_1_/G_0_ cells decreased slightly (Fig.2E). The bud S/G_2_/M population continued to expand, leveling out after six hours. This coincided with the appearance of the first G_1_/G_0_ cells contributing to the pEK. Also, at this stage, proliferation was specific to the tooth bud (Fig.S2D).

Shh and Fgf signaling have been suggested to induce proliferation in tooth buds (Hardcastle et al., 1998, Cobourne et al., 2009); however, inhibition of Shh signaling at the placode stage was reported to produce a flat wide molar bud with negligible effect on proliferation (Prochazka et al., 2015, Li et al., 2016b). This discrepancy could be explained by different effects of the signaling pathway on different molar cell populations. To substantiate this, we treated E11.5 cultures with cyclopamine to inhibit Shh signaling. Inhibition reduced bud invagination (Fig.S3A,C). This was concomitant with a reduced number of proliferating cells with expansion of G_1_/G_0_ phases adjacent to the IK. A few proliferating cells were still observed basally/peripherally (Fig.S3A,D). We next treated E11.5 cultures with an FGFR inhibitor (SU5402) to examine the role of FGF signaling. Inhibition of FGF signaling induced cell cycle exit somewhat later especially in basal cells, and less reduction in invagination and proliferation compared to cyclopamine treatment (Fig.S3B-D).

We, therefore, conclude that the molar IK is a functional signaling center that regulates proliferation in tooth bud invagination and growth. The molar bud is formed by localized cell proliferation, with the involvement of Shh and FGF signaling.

### IK ablation arrests progression of tooth development

To confirm that the IK drives molar bud growth and is necessary for progression of tooth development, we ablated the IK at different developmental stages by microsurgery and laser ablation. When the placode was microsurgically removed at E11.5 and the tissue cultured for 24h, no G_1_/G_0_ condensate was observed in the diastema and the epithelium remained flat (Fig.S4A,C). Microsurgical removal of the IK at E12.5 similarly arrested tooth growth, while development on the untreated side proceeded to bud stage with the emerging pEK present (Fig.S4B,C).

For a more targeted approach we next removed the IK G_1_/G_0_ cells at E11.5, E12.5 and E12.75 with laser ablation, followed by 24h culture, in K17-GFP and Fucci whole mount mandibles. Laser ablation of the IK G_1_/G_0_ cells in the early placode stage (E11.5) epithelium abrogated epithelial invagination and tooth development, while development in the control progressed normally (Fig.3A,C). Ablation at early bud stage E12.5 similarly completely arrested bud invagination and elongation and inhibited progression of tooth development (Fig.3B,C). Ablation somewhat later at E12.75 at a more developed bud stage also arrested growth; however, a small cluster of G_1_/G_0_ cells was observed in the bottom of bud facing the mesenchyme in the area where the pEK would emerge (Fig.3D). To observe if the arrested growth resulted from abrogated proliferation, we laser ablated the IK in the Fucci model. Correspondingly, ablation at E11.5 arrested invagination (Fig.3E) and this was accompanied with a loss in cell proliferation in the bud (Fig.3E,F). The persistence of S/G_2_/M cells in the adjacent oral epithelium confirmed good tissue health in non-ablated tissue (Fig.3E). Cell proliferation and consequently bud growth were similarly abrogated in E12.5 ablated molars (Fig.3G,H).

**Figure 3.**
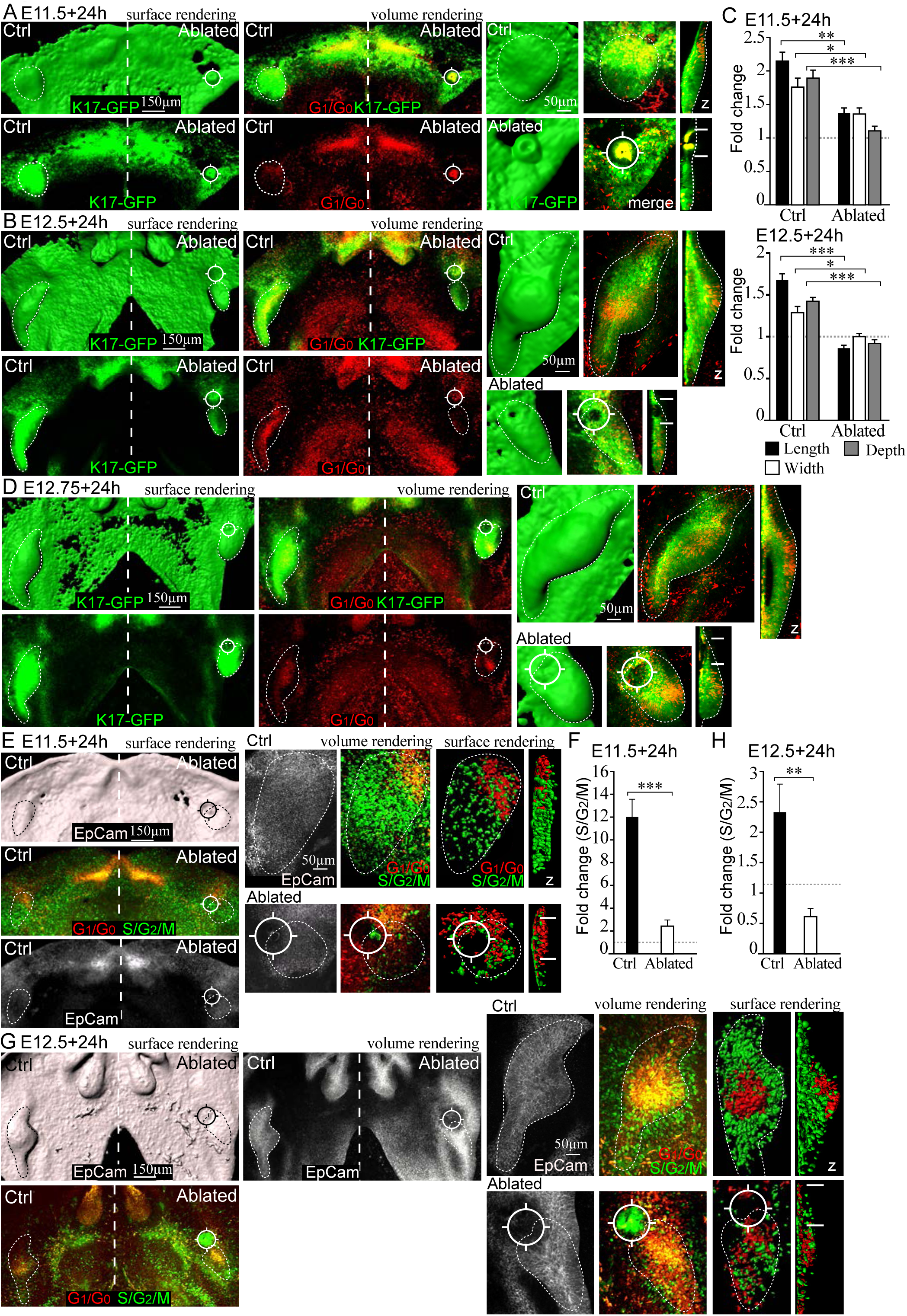
Laser ablation of the IK arrests molar bud growth. Confocal fluorescence images of whole mount explant cultures K17-GFP/ Fucci S/G_2_/M (green), Fucci G_1_/G_0_ (red), tooth placode/bud epithelium (dotted line), Hoechst (blue), IK (arrowhead) and pEK (asterisk). The IK was laser ablated (position marked with a viewfinder symbol ¤) at E11.5, E12.5 or E12.75 followed by 24h culturing. (A) Laser ablation of the IK G_1_/G_0_ cells in the early placode stage (E11.5) epithelium abrogated epithelial invagination and growth. In the control tooth invagination proceeded normally. (B) Early bud stage (E12.5) ablation completely arrested bud invagination and elongation. (C) Bud dimensions of ablated and control molars at E11.5+24h and E12.5+24h (fold change over E11.5, N_E11.5+24h_=9, N_E12.5+24h_=8, mean±SEM, Mann-Whitney U,* *p*≤*0*.*05, ** p*≤*0*.*01, *** p*≤*0*.*001*). (D) Ablation at E12.75 arrested bud growth. However, a cluster of G_1_/G_0_ cells emerged in the bottom of bud in the epithelium mesenchyme interface. (E) Laser ablation of the IK in the Fucci S/G_2_/M model at E11.5 resulted in loss of bud cell proliferation. S/G_2_/M cells present in the adjacent oral epithelium confirmed good tissue health. (F) Quantification of proliferating cells in E11.5+24h molars (N=8, mean±SEM Mann-Whitney U, *p*≤*0*.*001*). (G) Bud cell proliferation and consequently bud growth were similarly abrogated in E12.5 ablated molars. (H) Quantification of proliferating cells in E12.5+24h molars (fold change over E12.5, N=8, mean±SEM Mann-Whitney U, *p*≤*0*.*01*).

These experiments demonstrate that the molar IK is a functional signaling center that drives cell proliferation, thereby regulating tooth bud growth. The IK is, therefore, necessary for the progression of tooth development.

### The IK remains an integral part of the developing molar and does not contribute cells to the pEK

We next used whole-mount live tissue imaging to track individual cell movement in the different cell populations in placode and bud stage to dissect whether dynamical cell rearrangements contribute to molar bud formation. Signaling centers show canonical Wnt activity and we used the TCF/Lef:H2B-GFP reporter to visualize Wnt active cells together with the Fucci G_1_/G_0_ reporter to track signaling center cells. We further imaged the Fucci G_1_/G_0_ reporter with the S/G_2_/M reporter to follow the proliferating bud cell population.

Initially at E11.5, G_1_/G_0_ cells were distributed throughout the molar placode (Fig.4A,B,Mov.S3), similarly as observed in fixed samples (Fig.1A). More cells differentiated, entered G_1_/G_0_ cell cycle phase, and were redistributed mesio-lingually in the maturing placode. Tracking of IK cells showed that they moved toward the mesial front area of the bud remaining an integral part of the bud. In contrast, bud S/G_2_/M cells stayed mostly in place (Fig.4B). At E11.5 Wnt activity was seen throughout the dental lamina visualized by high TCF/Lef:H2B-GFP reporter fluorescence intensity (Wnt^Hi^) (Fig.4C,Mov.S4). The molar placode IK G_1_/G_0_ cells specifically localized to the peripheral border formed by dental lamina Wnt^Hi^ cells. Some overlap of G_1_/G_0_ and Wnt^Hi^ signal was seen but the G_1_/G_0_ cells mostly remained as a distinct subgroup (Fig.4C,D, Mov.S4). More G1 cells were recruited to the IK and they showed directional movement toward the dental lamina Wnt^Hi^ cells. The dental lamina Wnt^Hi^ cells and bud TCF/Lef:H2B-GFP+ cells remained non-motile (Fig.4C,D,Mov.S4).

**Figure 4.**
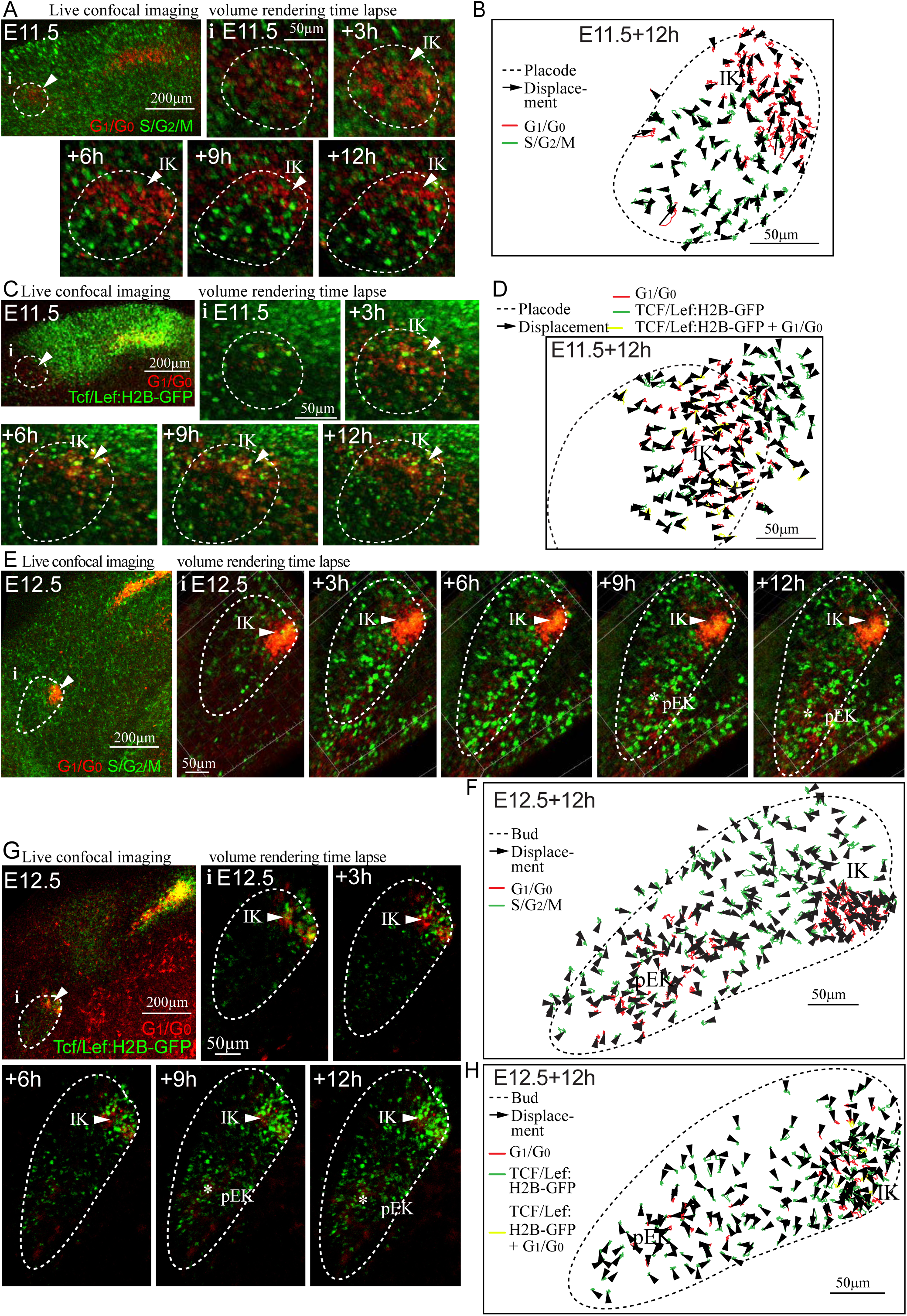
The IK remains an integral part of the tooth and does not contribute cells to the pEK that arises independently. (A) Still images of a Fucci G_1_/G_0_ (red) and S/G_2_/M (green) live tissue time lapse confocal imaging from placode stage E11.5+12h. (B) Tracks and displacement vectors of individual cells at E11.5+12h molar Fucci G_1_/G_0_, S/G_2_/M, placode epithelium border (dotted line). (C) Still images of a Fucci G_1_/G_0_ (red) and TCF/Lef:H2B-GFP (green) reporter time lapse E11.5+12h. (D) E11.5+12h cell tracks and displacement of individual Fucci G_1_/G_0_, TCF/Lef:H2B-GFP, cells and cells positive for both reporters. (E) Still images of a Fucci G_1_/G_0_ and S/G_2_/M reporter live imaging form early bud stage E12.5+12h. (F) E12.5+12h cell tracks and displacement of individual Fucci G_1_/G_0_ and S/G_2_/M cells (G) Still images of a Fucci G_1_/G_0_ and TCF/Lef:H2B-GFP live imaging E12.5+12h. (H) Cell tracks and displacement E12.5+12h Fucci G_1_/G_0_, TCF/Lef:H2B-GFP, and Fucci G_1_/G_0_+TCF/Lef:H2B-GFP.

Tracing cell movement in the molar IK and the emerging pEK from E12.5+12h showed that IK cells remained mesio-lingually close to the bud surface (Fig.4E,F,Mov.S5,Mov.S6). We did not detect contribution from either IK G_1_/G_0_ cells or Wnt^Hi^ to the pEK (Fig.4G,H,Mov.S6). The pEK arose deep in the bud, without clonal contribution from the IK (Fig.4G,H,Mov.S6). Also at this stage the S/G2/M cells showed little movement with no obvious orientation (Fig.4E,F,Mov.S5) and contribution of cells from oral epithelium, participating in bud growth, was not detected.

### Cell condensation and active directional cell migration drive molar IK maturation

Our live imaging experiments showed that IK cells reorganize dynamically during placode/bud maturation. To define the significance of this to IK maturation we quantified IK cell condensation and analyzed if the movements involve active cell migration.

We first measured cell density in EpCam stained fixed whole-mount samples. Initially, at E11.5, G_1_/G_0_ cells were more dispersed and at E12.5, they had condensed and retained this density until E13.5 (Fig.5A,B). Oral epithelial cells did not show a similar condensation. Quantification of cell density showed that condensation was specific to IK cells, with a significant increase in density from E11.5 to E12.5 compared to the oral epithelium (Fig.5B).

**Figure 5.**
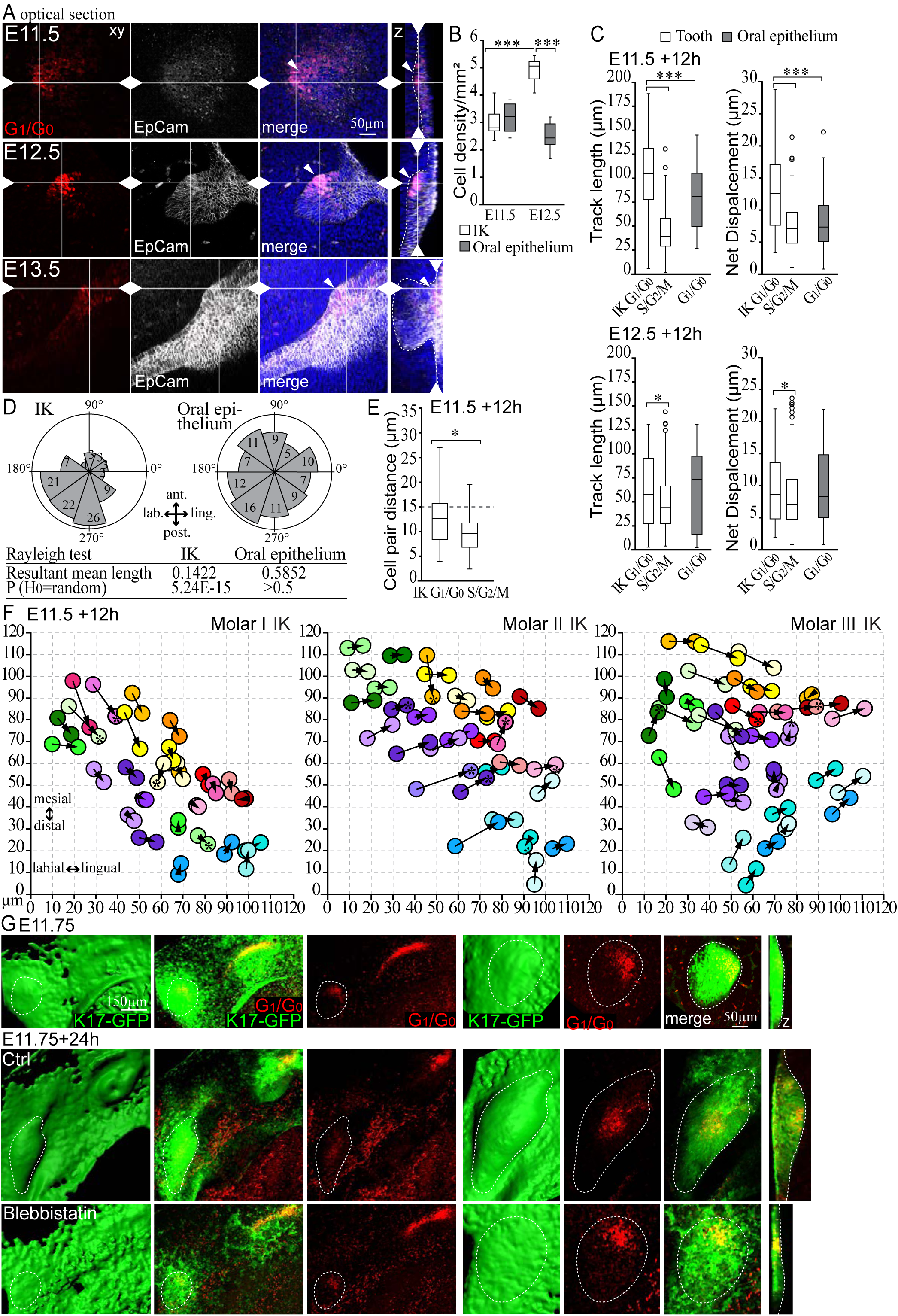
Cell condensation and active directional cell migration drive molar IK maturation. (A) Confocal fluorescence images of Fucci G_1_/G_0_ (red) molars, cell borders (EpCam, white, dotted line), Hoechst (blue). (B) Quantification of IK and oral epithelial cell density (N_placode/bud_=N_oral_=10, Mann-Whitney U, *p*≤0.001). (C) Quantification cell of track length and net displacement in molar placode/bud and oral epithelium at E11.5+12h (N_G1/G0 IK_=149, N_G1/G0 oral_=45, N_S/G2/M_=146, Mann-Whitney U, *p*≤0.001***) and E12.5+12 h (N_G1/G0 IK_=90, N_G1/G0 oral_=51, N_S/G2/M_=317, Mann-Whitney U, *p*≤0.05). (D) Quantification of molar IK and oral epithelial cell movement angles E11.5+12h. (N_IK cells_=N_oral epithelial cells_=95, Rayleigh test: H_0_=random, IK *p*≤0.001, oral *p*>0.05). (E) Pairwise comparison of molar IK G_1_/G_0_ and bud S/G_2_M cell positions (N_pairs_=40, Mann-Whitney U test, *p*≤*0*.*05)*. (F) Tracing of groups of cells at E11.5 that resided close to each other (respectively color coded) and positions of the individual cells 12h later. Cells switching neighbors (asterisk). (G) Confocal fluorescence whole mount images of Fucci G_1_/G_0_ (red) and K17-GFP (green) reporter. E11.75 cultures were treated at the time of most active IK G_1_/G_0_ cell movement with blebbistatin for 24h to inhibit actomyosin based cell motility. Blebbistatin treatment repressed IK G_1_/G_0_ cell condensation and arrested bud morphogenesis.

To study if IK condensation is achieved through active cell migration, we followed the movement of individual cells by live imaging at E11.5/E12.5+12h. Tracking showed active migration of the molar IK G_1_/G_0_ cells at both time points. We quantified the overall track length and net displacement in the different cell populations, and at E11.5+12h, a significant difference in IK G_1_/G_0_ cells was observed: They migrated more compared to both oral epithelial G_1_/G_0_ and tooth bud S/G_2_/M cells (Fig.5C). At E12.5+12 h, IK G_1_/G_0_ cells still showed a longer mean track length in the bud compared to S/G2/M cells (Fig.5C). Quantification of IK G_1_/G_0_ cell displacement angles at E11.5+12h showed a distinct orientation towards the mesio-lingual side of the forming placode/bud, whereas oral epithelial cells showed a random orientation (Fig.5D). We confirmed active migration by following pairs of IK G_1_/G_0_ and bud S/G_2_/M cells that were initially in close proximity (distance≤15µm). The pairwise comparison revealed that while many IK G_1_/G_0_ cells remained neighbors, 30% of cells switched their partners; most S/G_2_/M pairs remained neighbors (Fig.5E). Tracing groups of cells that were initially, at E11.5, located next to each other in different areas of the IK, and defining cell positions 12h later showed that some cells remained close to their original neighbors but several cells ended up with a different group. (Fig.5F). Pharmacological inhibition of acto-myosin based motility, with the inhibitor blebbistatin, repressed IK G_1_/G_0_ condensation and abrogated progression of tooth development (Fig.5G).

### Dynamics between Wnt^HI^ and *Shh* cell populations regulate the maturation and maintenance of the IK

Our live imaging analysis suggested that TCF/Lef:H2B-GFP reporter expressing Wnt^Hi^ cells were closely juxtaposed to *Shh* expressing G_1_/G_0_ IK cells but they seemed to comprise two different cell populations that remained in close contact with each other through bud development (Fig.6,Mov.S4, Mov.S6,Fig.S1E). The Shh pathway is an important modulator for Wnt signaling for several stages of tooth development. Studies in mouse mutants have implied that Shh is a downstream target of Wnts and also an inhibitor of Wnt signaling via a negative feedback loop (Sarkar, et al., 2000; Sarkar and Sharpe, 1999). We next investigated the behavioral dynamics and the molecular identity of the two populations.

**Figure 6.**
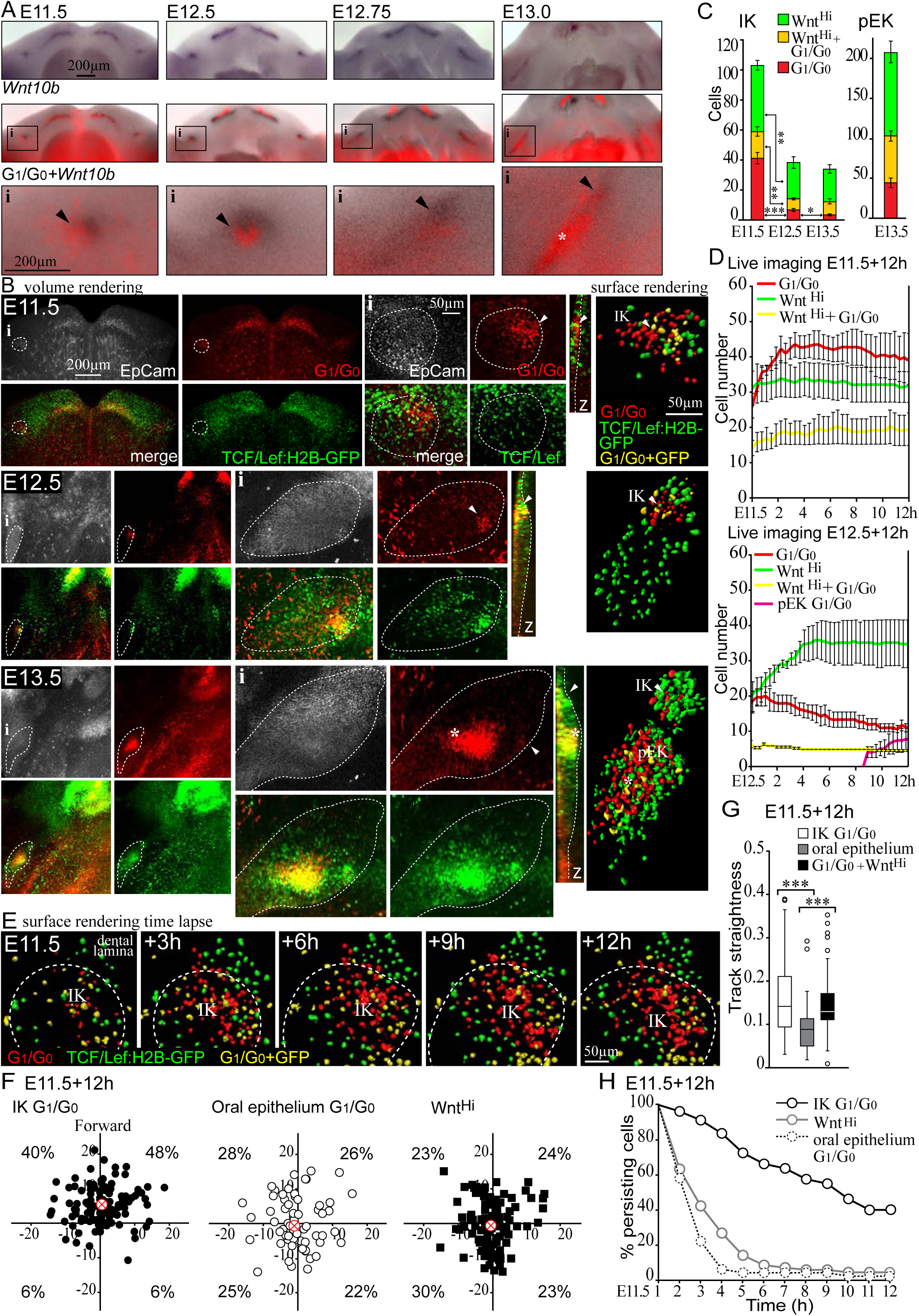
Dynamics between Wnt^HI^ and *Shh* cell populations regulate the maturation and maintenance of the IK. (A) Fucci G_1_/G_0_ fluorescence images overlaid with whole-mount DIG *in situ* hybridization for *Wnt10b*. (B) Single cell resolution analysis of Fucci G_1_/G_0_ (red) and TCF/Lef:H2B-GFP (green), reporter for active Wnt/β-catenin signaling, patterns. EpCam (white, dotted line), IK (arrowhead) and pEK (asterisk). At E11.5, TCF/Lef:H2B-GFP high intensity (Wnt^Hi^) was present in the dental lamina and the molar IK G_1_/G_0_ condensate was located next to a border of Wnt^Hi^ cells. At E12.5, IK G_1_/G_0_ only cells were surrounded by Wnt^Hi^ cells. At E13.5, G_1_/G_0_ and Wnt^Hi^ cells were present in the presumptive pEK region. (C) Quantification number of cells in Wnt^Hi^, G_1_/G_0_ and double positive cell populations in fixed samples (N_E11.5_=413; N_E12.5_=192; E13.5 N_IK_=136 N_pEK_=831, mean±SEM, non-parametric Student’s t-test, *p*≤0.05*, *p*≤0.01**, *p*≤0.001***). (D) Quantification of cell number in each population from live tissue imaging E11.5+12h and E12.5+12h (N_E11.5+12h_ =N_E12.5+12h_=3, mean±SEM). (E) Surface rendering still images of E11.5+12h molar time-lapse, Fucci G_1_/G_0_, TCF/Lef:H2B-GFP, and double cells. At E11.5+3h onwards the G_1_/G_0_ cell population started differentiating closely juxtaposed to Wnt^Hi^ cells with increasing number of cells transitioning to G_1_/G_0_. (F) Track end point analysis (median: red cross, N_IK G1/G0_=128, N_oral G1/G0_=63, N_WntHi_=104). Molar IK G_1_/G_0_ cells showed preferential distribution toward the Wnt^Hi^ region (forward). (G) Track straightness in molar IK G_1_/G_0_, oral epithelial G_1_/G_0_ and TCF/Lef:H2B-GFP+ cell populations (N_IK G1/G0_=128, N_oral G1/G0_=42, N_WntHi_ =67, Mann-Whitney U, *p*≤0.001). (E) Decay of cellular persistence (N_IK G1/G0_=80, N_oral G1/G0_=50, N_WntHi_=71).

*In situ* hybridization analysis of Fucci specimens revealed that the IK G_1_/G_0_ cells colocalized with *Shh* (Fig.S1E). A canonical Wnt, *Wnt10b*, has previously reported expression in the placode (Liu, et al., 2008). When we did a *Wnt10b* hybridization in the Fucci G_1_/G_0_ reporter, at E11.5; however, *Wnt10b* expression was also detected, partially overlapping, but predominantly anterior to the G_1_/G_0_ focus (Fig.6D). By E12.5, the dense IK G_1_/G_0_ colocalized with the *Shh* signal, whereas the *Wnt10b* expression covered a larger area surrounding the IK G_1_/G_0_ condensate. By E12.75, the diffuse *Wnt10b* expression continued to reside in a wider area in the molar mesio-lingual tip. At E13.0, the G_1_/G_0_ area was barely discernible and *Shh* and *Wnt10b* were downregulated. Concomitantly, G_1_/G_0_, *Shh*, and *Wnt10b* expression appeared in the emerging pEK (Fig.S1E, Fig.6A,D). There was a complete spatial correlation with *Shh* and G_1_/G_0_ signal throughout early molar morphogenesis, but *Wnt10b* expression was seen in the area juxtaposing the G_1_/G_0_ focus anteriorly.

High resolution analysis of G_1_/G_0_ and TCF/Lef:H2B-GFP patterns showed that the *Shh*-G_1_/G_0_ cell population initiated at E11.5 was closely juxtaposed to Wnt^Hi^ cells (Fig.6B,Mov.S4). By E12.5, Wnt^Hi^ cells surrounded the *Shh*-G_1_/G_0_ cells and TCF/Lef:H2B-GFP+ cells were scarcely distributed in the growing bud prior to pEK appearance (Fig.6B,Mov.S6). At E13.5, Wnt^Hi^ cells were present in the pEK with G_1_/G_0_ cells distributed more centrally (Fig.6B). Quantification of *Shh*-G_1_/G_0_ and Wnt^Hi^ cell populations in fixed samples showed a decrease at E12.5 in *Shh*-G_1_/G_0_ cell number, concomitant with TCF/Lef:H2B-GFP downregulation and appearance of apoptosis specifically in the IK cells (Fig.6C, Fig.S5A,B) consistent with canonical Wnt signaling activity and *Shh* expression in the maintenance of the IK. To more closely examine this dynamic, we quantified the cell *Shh*-G_1_/G_0_ and Wnt^Hi^ populations with live imaging in E11.5 and E12.5+12h molars. Quantification at E11.5+12h showed that the number of Wnt^Hi^ cells remained stable for the 12h follow-up; in contrast, the *Shh*-G_1_/G_0_ cell population increased by 1.5-fold (Fig.6D). Analysis of E12.5+12h cells, showed an increase in Wnt^Hi^ cells, throughout the bud, reaching a plateau after 4h; IK *Shh*-G_1_/G_0_ cells showed a constant decrease, and G_1_/G_0_ cells appeared in the pEK from 9h onward (Fig.6D).

Analysis of individual contributing cell populations in the initiation of the molar placode showed a border region with an accumulation of Wnt^Hi^-*Wnt10b* cells in the dental lamina and the G_1_/G_0_-*Shh* cells started differentiating closely juxtaposed to this region (Fig.6E). Analysis of cell movement showed differential patterns in the Wnt^Hi^-*Wnt10b* and G_1_/G_0_-*Shh* cell populations: track end point analysis showed specific preferential movement of G_1_/G_0_-*Shh* IK cells toward the dental lamina Wnt^Hi^ cells (Fig.6F) with high straightness (Fig.6G) and high directional persistence in the G_1_/G_0_-S*hh* IK cell compared to oral epithelial G_1_/G_0_ and dental lamina Wnt^Hi^ cells (Fig.6H).

The differential distribution and cellular behaviors of Wnt^Hi^ and *Shh*-G_1_/G_0_ cells in the molar signaling centers suggests that they act in concert to initiate signaling center cell differentiation in the very early stages. The boundary between the two cell populations defines the position of the emerging molar IK and orients the directional migration pattern for condensation. Further, decreased Wnt signaling resulted in *Shh* downregulation and IK clearance.

### Modulation of canonical Wnt signaling affects IK cell dynamics and tooth bud shape

The cell movement data suggested the presence of a chemotactic gradient from the dental lamina Wnt^Hi^-*Wnt10b* cells directing the movement and condensation of the G_1_/G_0_-*Shh* cell population in molars. *Wnt10b* has been implicated as a paracrine chemotactic factor in epithelial cancer contexts (Chen, et al., 2017; Aprelikova, et al., 2013). To explore if this dynamic occurs in developing molars, we modulated canonical Wnt signaling levels by stimulation with Wnt3a or a Wnt10b releasing bead, and by inhibition with a Wnt antagonist, XAV939 that acts via stimulation of β-catenin degradation and stabilization of axin. E11.5 explants were treated either with Wnt3a/XAV939 in the growth medium for 24h or by placing a recombinant Wnt10b soaked/control bead next to the placode at E11.5 and followed the explants for 16h.

We used K17-GFP to visualize the shape of the epithelium and Fucci G_1_/G_0_ for IK cell distribution in the developing placode/bud. Stimulation with Wnt3a resulted in a flat bud compared to control (Fig.7A,B), with persisting number of G_1_/G_0_ IK cells spread out throughout the invagination (Fig.7A,C). Inhibition of active Wnt signaling with XAV939 resulted in a complete loss of G_1_/G_0_ condensate together with a loss of invagination (Fig.7A). To study if lack of IK condensation and the loss of invagination, with Wnt modulation, was caused by lack of bud cell proliferation, we treated Fucci G_1_/G_0_; S/G_2_/M mandibles with Wnt3a or XAV939. Stimulation with Wnt3a resulted in lack of IK condensation followed by a drastic loss of cell proliferation in the bud (Fig.7D,E). Inhibition with XAV939 resulted in the loss of the G_1_/G_0_ IK condensate and absence of proliferation and invagination (Fig.7D).

**Figure 7.**
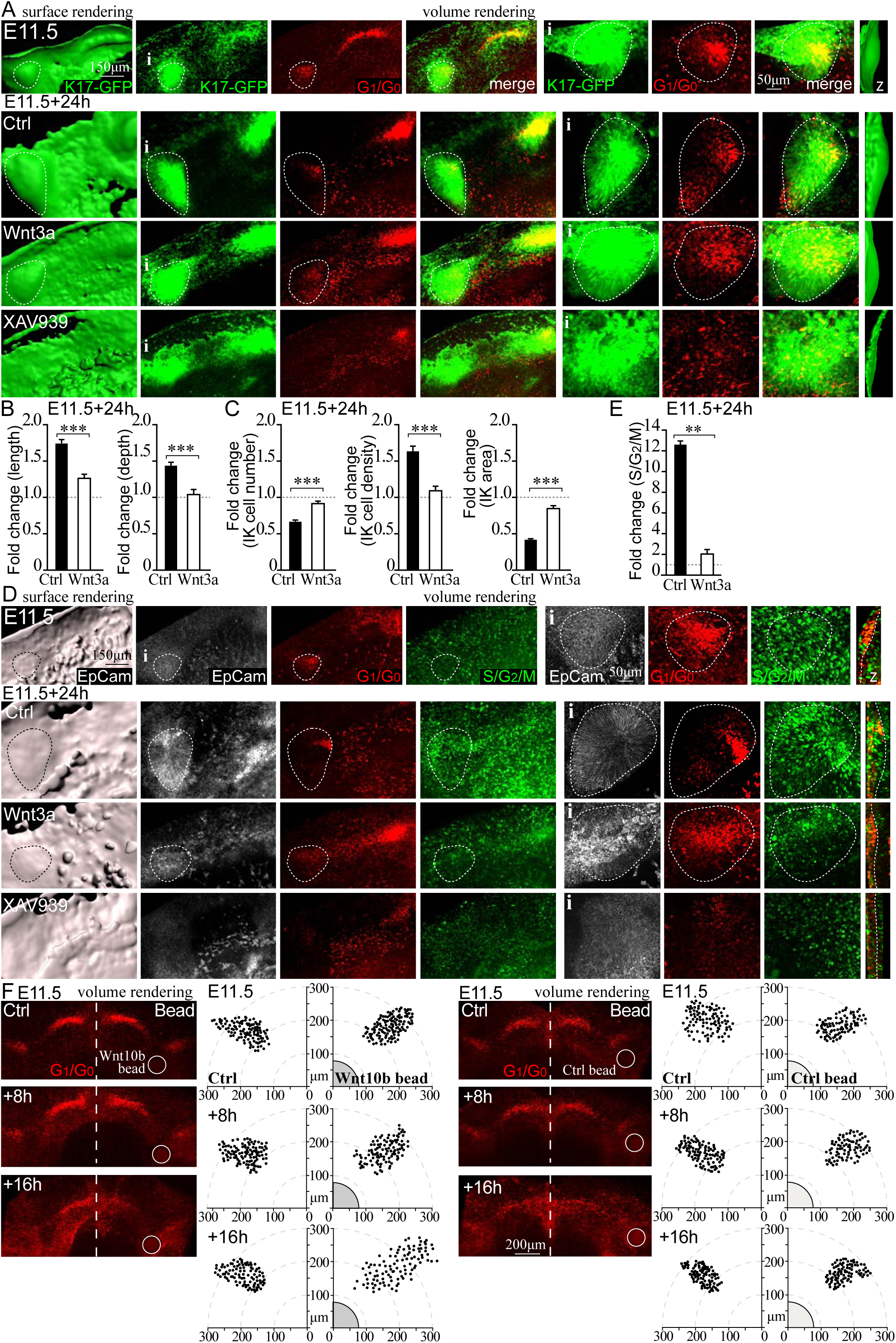
Modulation of canonical Wnt signaling affects IK cell dynamics and molar bud shape. (A) Confocal fluorescence images of explants cultures K17-GFP (green), Fucci G_1_/G_0_ (red), tooth placode/bud epithelium (dotted line). Canonical Wnt signaling levels were modulated from placode stage E11.5+24h by stimulation with Wnt3a, of inhibition with XAV939. Stimulation resulted in a flat bud with persisting G_1_/G_0_ IK cells throughout the invagination. Inhibition lead to a complete loss of the G_1_/G_0_ condensate and abrogated invagination. (B) Quantification of bud dimensions in Wnt3a stimulated and control cultures (fold change over E11.5, N_ctrl_=12, N_Wnt3a_=10, mean±SEM, Mann-Whitney U, *p*≤*0*.*001*). (C) Quantification of IK cell number, density and area in Wnt3a stimulated and control cultures (fold change over E11.5, N_ctrl_=12, N_Wnt3a_=10, mean±SEM, Mann-Whitney U, *p*≤*0*.*001*). (D) Stimulation with Wnt3a in Fucci G_1_/G_0_; S/G_2_/M cultures resulted in lack of IK condensation followed by a drastic loss of cell proliferation in the bud. Inhibition with XAV939 resulted in the loss of the G_1_/G_0_ IK condensate and absence of proliferation and invagination (E) Quantification of cell proliferation in Wnt3a stimulated and control cultures (fold change over E11.5, N_ctrl_=7, N_Wnt3a_=8, mean±SEM, Mann-Whitney U, *p*≤*0*.*01*). (F) A Wnt10b recombinant protein releasing or control bead was placed close to the IK (visualized with Fucci G_1_/G_0_ reporter) distally on the lingual side of the placode and the explants were imaged at E11.5, after 8h, and 16h. Morphogenesis and IK condensation proceeded normally in control buds without beads and with a control bead. The Wnt10b bead showed a loss in condensation of the G_1_/G_0_ IK cells. Instead the G_1_/G_0_ IK cells were spread out toward the Wnt10b bead.

To dissect the role of *Wnt10b* on IK condensation, we placed a Wnt10b releasing or control bead close to the IK distally on the lingual side of the placode and imaged the explants at E11.5, after 8h, and 16h. While morphogenesis and IK condensation proceeded normally in both the control bud and the bud with the control bead, the bud with the Wnt10b bead showed a loss in condensation of the G_1_/G_0_ IK cells (Fig.7F). Instead, the G_1_/G_0_ IK cells were spread out toward the Wnt10b bead. Measurement of bud dimensions showed a change in bud shape, e.g. decrease in bud elongation, in Wnt10b bead buds together with a decrease in G_1_/G_0_ IK cell density (Fig.S6A). The changes in IK cell distribution and bud shape were also accompanied with decrease in proliferation (Fig.S6B, C).

## Discussion

The reiterative genetic regulation of tooth development via signaling centers is conserved across tooth types, but it is less understood how it is interpreted into different cellular behaviors to regulate tooth shape and size. In the present study we have identified a molar IK signaling center that is necessary for the progression of tooth development in the early stages of mammalian tooth morphogenesis: We show with live imaging, 4D whole-mount analyses, and functional ablation studies that the IK arises in the placode and is a functional signaling center that drives proliferative growth prior to the successive enamel knots. Molar IK cell dynamics displays the hallmarks of ectodermal signaling centers: cell cycle exit and condensation, and silencing through apoptosis. Cell cycle exit coupled to active condensation, takes place not only in teeth (current study, Ahtiainen et al. 2016), but also in hair placodes (Ahtiainen, et al., 2014). Condensation of the IK via active cell movement is necessary for progression of tooth budding, as inhibition of actomyosin contractility and modulation of condensation guiding Wnt signaling levels, compromised condensation and the function of the signaling center. Cell condensation could be a universal mechanism to both trigger the ectodermal signaling center differentiation and cell cycle exit by contact inhibition of cell proliferation and eventually regulate timing of signaling center silencing by initiating mechanical crowding induced apoptosis.

Tissue recombination studies have shown that the instructive potential in the tooth first resides in the epithelium and shifts only later to the mesenchyme (Lumsden, 1988; Mina and Kollar, 1987). EKs require inductive signals from the mesenchyme, but it is plausible that the IK inducing signal comes from planar epithelial signaling. Wnts*7b/3* and Shh have a mutually exclusive expression already at E10.5 in the presumptive oral and dental ectoderm (Sarkar, et al., 2000; Sarkar and Sharpe, 1999) so it appears that different Wnt expression patterns and *Shh* determine the ectodermal boundaries of competence at a very early stage. *Shh* is possibly a downstream target of Wnts and also a negative feedback inhibitor. Spatial inhibition of *Wnt10b* by *Shh* has been reported in teeth: Shh coated beads repressed *Wnt10b* but no other epithelial markers (Dassule and McMahon, 1998). Constitutive activation of epithelial Wnt/β-catenin, somewhat later from E12.5, induced multiple patches of signaling center markers, including *Shh* and *Wnt10b* at E13-E14, and ectopic teeth (Liu, et al., 2008; Jarvinen, et al., 2006). Our work evidences that *Wnt10b* and *Shh* are differentially expressed during molar initiation and that these cell populations remain functionally separate. However, close interaction between the G_1_/G_0_-*Shh* and Wnt^Hi^-*Wnt10b* expressing cells is crucial in the positioning and maintenance of the molar IK. Wnt10b has been implicated as a paracrine chemotactic factor in cancer contexts (Chen, et al., 2017; Aprelikova, et al., 2013). The migration of G_1_/G_0_-*Shh* IK cells toward the canonical Wnt gradient and specific area of endogenous *Wnt10b* expression and distribution of IK cells toward exogenous source of recombinant Wnt10b, suggest that *Wnt10b* carries an instructive role in signaling center condensation.

We show that molar invagination and growth take place through cell proliferation in both basal and suprabasal bud cell populations, driven by the non-proliferative IK. Several signaling pathways regulate behaviors in these cell populations, but the role of Hh signaling in this context has been debated. Shh has been interpreted to be a primary inducer of proliferation in some experimental settings, whereas other studies suggest a role in bud cell rearrangement (Li, et al., 2016b; Prochazka, et al., 2015; Cobourne, et al., 2009; Hardcastle, et al., 1998). *Shh* expression is a hallmark of signaling centers, and while autocrine signaling cannot be ruled out, most of the responsive cells seem to reside elsewhere: at later stages, from E14.5 onwards, the pEK expresses *Shh* but receptor *Ptch* and downstream targets *Gli1/2/3* are expressed in the mesenchyme (Hardcastle, et al., 1998; Vaahtokari, et al., 1996a). In early stages from E11.0, the transducer *Smo* is ubiquitously expressed, whereas *Ptch1* and *Gli1* are mostly in the mesenchyme (Dassule and McMahon, 1998; Hardcastle, et al., 1998). Interestingly, by E12.0, the expression of both *Ptch1* and *Gli1/2* have been reported in the emerging epithelial bud. Notably, in our analyses, proliferation coincided with this. The inhibition of Shh signaling at the placode stage resulted in loss of proliferation in the bud. Thus, it seems that in the IK, *Shh* expressing cells are different from Shh responsive cells and signaling can induce proliferation. In agreement, early findings from conditional *Shh* mutants show smaller teeth and posteriorly misplaced buds (Dassule, et al., 2000; Dassule and McMahon, 1998). In the limb bud, Shh has been reported to affect proliferation both directly and indirectly via induction of Fgfs in the AER (Prykhozhij and Neumann, 2008; Towers, et al., 2008). In the tooth, other factors downstream of the initial placodal inducer Shh, such as Fgfs, likely contribute to bud invagination via proliferation mainly in basal cells, and possibly concurrent with stratification (current study, (Li, et al., 2016b).

Expression levels of *Shh* may specifically affect signaling center identity, differentiation, and maintenance. *Shh* overexpression arrests development at the bud stage due to lack of proliferation, and multiple superficial invaginations are induced in the epithelium but still fail to develop further (Cobourne, et al., 2009). It is tempting to hypothesize that exogenous *Shh* expression would drive cells into abnormal cell cycle exit and/or a change in fate into a signaling center. Signaling center maintenance may also be associated with *Shh* expression levels. Shh has been shown to be protective of early apoptosis in the tooth (Cobourne, et al., 2001). Apoptosis is a mechanism used to silence signaling centers in teeth, the AER of the limb, and in embryonic brain development (Nonomura, et al., 2013; Matalova, et al., 2004; Vaahtokari, et al., 1996b). The interplay between Wnt^Hi^ and *Shh*+ cells may serve as a feedback mechanism regulating the timing of apoptosis in the IK.

We demonstrate that the pEK in the molar is formed de novo without clonal contribution from the IK, and that the IK is apoptotically silenced upon pEK appearance. This differs mechanistically from signaling centers later in molar development, where the pEK contributes cells to sEKs (Du, et al., 2017). The development of teeth is conserved, however, in being driven by the iterative use of signaling centers. We have shown functionally here that the progression of early molar morphogenesis is dependent on the IK signaling center that arises in the placode and exhibits many hallmarks of ectodermal signaling centers. What differentiates the IK from the later signaling centers on a transcriptomic level will be of special interest to future studies.

## Acknowledgments

We thank Frederic Michon and Jukka Jernvall for critical reading of the manuscript. Imaging was done at the Light Microscopy Unit/Institute of Biotechnology, University of Helsinki. The work was financially supported by the Academy of Finland, the Sigrid Jusélius Foundation, Finnish Cultural Foundation and Helsinki Institute of Life Sciences.

The authors declare no competing financial interests.

## Author Contributions

Conceptualization L.A. and I.M.; Methodology, I.M., M.N., L.A.; Investigation I.M., M.N., L.A; Writing Original draft, L.A., I.M. and J.M-V; Writing, Review and Editing L.A., I.M., J.M-V and M.N.; Funding Acquisition L.A.; Supervision L.A.

## Materials and Methods

### Animals, tissues preparation and culture treatments

All mouse studies were approved by the National Animal Experiment Board. Transgenic mouse reporter lines: fluorescent cell cycle indicator (Fucci) mice express a nuclear red fluorescent reporter in G_1_/G_0_ phase (Cdt1-mKO) and green fluorescent reporter in S/G_2_/M phases (Gem-mAZ) (Sakaue-Sawano, et al., 2008). Shh^GFPCre^ mice (#005622, Jackson Laboratories) express GFP consistent with endogenous *Shh* locus marking signaling centers (Harfe, et al., 2004), K17-GFP mice visualize the tooth epithelium (#023965, Jackson Laboratories). TCF/Lef:H2B-GFP mice, are indicators of Wnt/β-catenin signaling, containing several copies of TCF/Lef1 DNA binding sites driving expression of the H2B-EGFP fusion protein (#013752; Jackson Laboratories); FGF20^βGal^ mice have an Fgf20-β-galactosidase (βGal) knock-in allele (Huh, et al., 2012). Embryos were staged according to limb morphological criteria; vaginal plug day was embryonic day (E)0.5 (Martin, 1990).

Embryonic mandibles were dissected at E11.5-E13.5 and whole mount explants where fixed from 2h to overnight in 4% PFA or cultured in a Trowell-type tissue culture as described previously (Narhi and Thesleff, 2010). For live imaging experiments tissues where maintained in D-MEM/F12 without phenol red and supplemented with 50U/ml penicillin, 50µg/ml streptomycin, 10% FCS and HEPES 15mM (Gibco). For inhibitor/activator treatments samples were dissected at E11.5 or E11.75, and vehicle or smoothened inhibitor cyclopamine was applied for inhibition of Shh signaling (50µM; Sigma-Aldrich), blebbistatin to inhibit actomyosin mediated cell motility (100µM; Sigma-Aldrich), pan-FGFR inhibitor SU5420 (20µM; Calbiochem), recombinant Wnt3a (10ng/ml; R&D Systems) for stimulation and XAV939 (10μM; Tocris) to inhibit canonical Wnt signaling was added to the growth medium for 24h. For bead implantation, heparin acrylic beads (MCLAB) were incubated with 100μg/ml recombinant human Wnt10b protein (0.1mg/ml;R&D Systems) at 37°C for 30 minutes. Control beads were soaked with similar concentrations of BSA under the same conditions. Protein-soaked beads were stored at 4°C and used within one week. Beads were applied on tissue explant cultures at E11.5 distally on the lingual side of the placode and tissues were imaged at the start point, after 8 and 16h to ensure good tissue condition.

### Whole mount immunofluorescence, fluorescence microscopy and *in situ* hybridization

For whole mount immunofluorescence staining fixed tissues were permeabilized with 0.5% TritonX-100 for 2h RT and washed with PBS. Unspecific staining was blocked by incubation in 5% normal donkey/goat serum, 0.3% BSA, 0.1% TritonX-100 in PBS 1h RT. Tissues were incubated overnight in +4°C with the primary antibody rat polyclonal anti-mouse CD326 (EpCam, 1:1000, Pharmingen), rabbit polyclonal βGal (1:400, MP Biomedicals), rabbit polyclonal cleaved caspase 3 (1:400, Cell Signaling Technologies) or goat polyclonal sonic hedgehog antibody (1:100, R&D Systems) and detected with Alexa Fluor-488, Alexa Fluor-568 or Alexa Fluor-647 conjugated secondary antibodies (1:500, BD and Invitrogen) and nuclei were stained with Hoechst 33342. Tissues were mounted with Vectashield (Vector Laboratories) and imaged with either Leica Biosystems TCS SP5 microscope and HC PL APO 10×/0.4 (air), HCX PL APO 20×/0.7 Imm Corr (water, glycerol, oil) Lbd.bl and HCX APO 63×/1.30 Corr (glycerol) CS 21 objectives or Zeiss LSM700 microscope and HC PL APO 10×/0.45 (air) and LD LCI PL APO 25×/0.8 Imm Corr (water, glycerol, oil) objectives. For analysis of TCF/Lef:H2B-GFP signal intensities, the cutoff value for high and low expressing cells was adjusted according to overall signal intensity in each sample. All results represent at least three independent experiments.

For combined fluorescence and whole mount *in situ* hybridization analyses: Fluorescent imaging for fixed Fucci G_1_/G_0_ reporter whole mount mandibles was first done with a Zeiss SteREO Lumar.V12 microscope, NeoLumar S 0.8×/WD 80-mm objective, and Zeiss Axiocam MRm3 CCD camera. The samples where then subjected to whole mount *in situ* hybridization with digoxigenin-labeled probes specific for *Shh* or *Wnt10b* performed as described previously (Shirokova, et al., 2013; Fliniaux, et al., 2008; Wang and Shackleford, 1996). Imaging of the hybridization signal was done with the same Zeiss Lumar microscope and Zeiss AxioCam ICc1 CCD camera and images were transposed.

### Fluorescence confocal microscopy and time-lapse imaging and laser ablation

For 3D time-lapse imaging dissected tissues were allowed to recover for a minimum of 2h prior to imaging. The explants were imaged as described previously (Ahtiainen, et al., 2016; Ahtiainen, et al., 2014) with an upright Leica Biosystems TCS SP5 microscope, HC PL APO 10×/0.4 (air) objective in a trowel-type culture setup. Z-stacks of 3µm optical sections were acquired at 20min intervals. Good tissue health was confirmed by lack of pyknotic nuclei and frequency of mitoses in every acquired z-stack. For determination of cell cycle status and cell quantification, only cells that were distinctly identified as either G_1_/G_0_ or S/G_2_/M were scored. For Wnt/β-catenin signaling activity TCF/Lef:H2B-GFP cells were scored individually for median fluorescence intensity in each nucleus and presence of G_1_/G_0_ signal. All results represent at least three independent experiments.

Laser ablations were done with an upright Leica Biosystems TCS SP5 microscope, HC PL APO 10×/0.4 (air) objective and a tunable Ti:Sapphire pulsed IR laser (Spectra Physics, MaiTai, tunable range 690-1040nm) at room temperature with an excitation wavelength 800nm and 2.95W of laser power (100%) for 3-10 seconds. The pulse was targeted to the IK, visualized with the Fucci reporter, using zoom factor 20-40x. Efficiency of ablation was verified by acquiring confocal fluorescence Z-stacks of the sample immediately after ablation. After 24h of culture tissues were fixated, Fucci cell cycle reporter samples were immunofluorescence stained with EpCam to visualize the epithelium, and all samples were stained with Hoechst 33342 to visualize nuclei. Samples were imaged with the Zeiss LSM700 microscope, with HC PL APO 10×/0.45 (air) and LD LCI PL APO 25×/0.8 Imm Corr (glycerol) objectives. Specificity of ablation was verified by the absence of Fucci G_1_/G_0_ phase (Cdt1-mKO) positive cells in the IK region and ablation of only the epithelial compartment visualized with the K17-GFP reporter and Hoechst staining. Good tissue health of the adjacent, non-ablated tissue, was confirmed by lack of pyknotic nuclei and presence of normal cell proliferation patterns with the Fucci reporter.

### Quantitative and statistical analyses of experimental data

Analyses of images and quantitative measurements were done with Imaris 9.0.1 (Bitplane) and ImageJ software. Images were processed for presentation with Photoshop CC and Illustrator CC software (Adobe Systems). Statistical analysis and further graphing were done with PAST (http://folk.uio.no/ohammer/past/; Hammer et al., 2001), and SPSS Statistics (IBM) software.

Measurements were done from whole mount volume renderings of confocal optical Z-stacks. For quantifying cell density individual cell borders were visualized and traced in 3D with the EpCam staining in whole-mount tissues. Cell densities were quantified by masking a volume in tooth epithelium/equal volume in the oral epithelium, and defined as areas occupied by the cell (selecting a cross section in 3D view in the middle of the cell). Box-and-whiskers plots represent minimum, 25th percentile, median, 75th percentile, and maximum values for each dataset. Differences between groups were assessed with the Mann-Whitney U test.

All cell movement, follow up and division analyses were done from stereoscopic 3D renderings, allowing exact localization of cells in three dimensions, and in time. Individual cell track length and net displacement were measured in signaling center and oral epithelial cells. The distribution of cell trajectory displacement angles was analyzed with the Rayleigh test (H0=random, *p*>0.05). For IK G_1_/G_0_ group and pairwise cell trajectory analysis tissues were live imaged E11.5+12h. G_1_/G_0_ cells where divided into neighboring groups in their original position and traced to the end position. Cells within close proximity of each other (≤15µm) were analyzed in pairs. For the analysis on decay of cellular persistence in directional migration, we first determined the angle of cell migration during the first hour of observation for the initial orientation of the cells. At each following time point, cells that had not yet turned > ±90° from their starting angle, were considered directionally persistent.

## Supplemental Figure Legends

**Related to Figure 1**.

**Figure S1.**
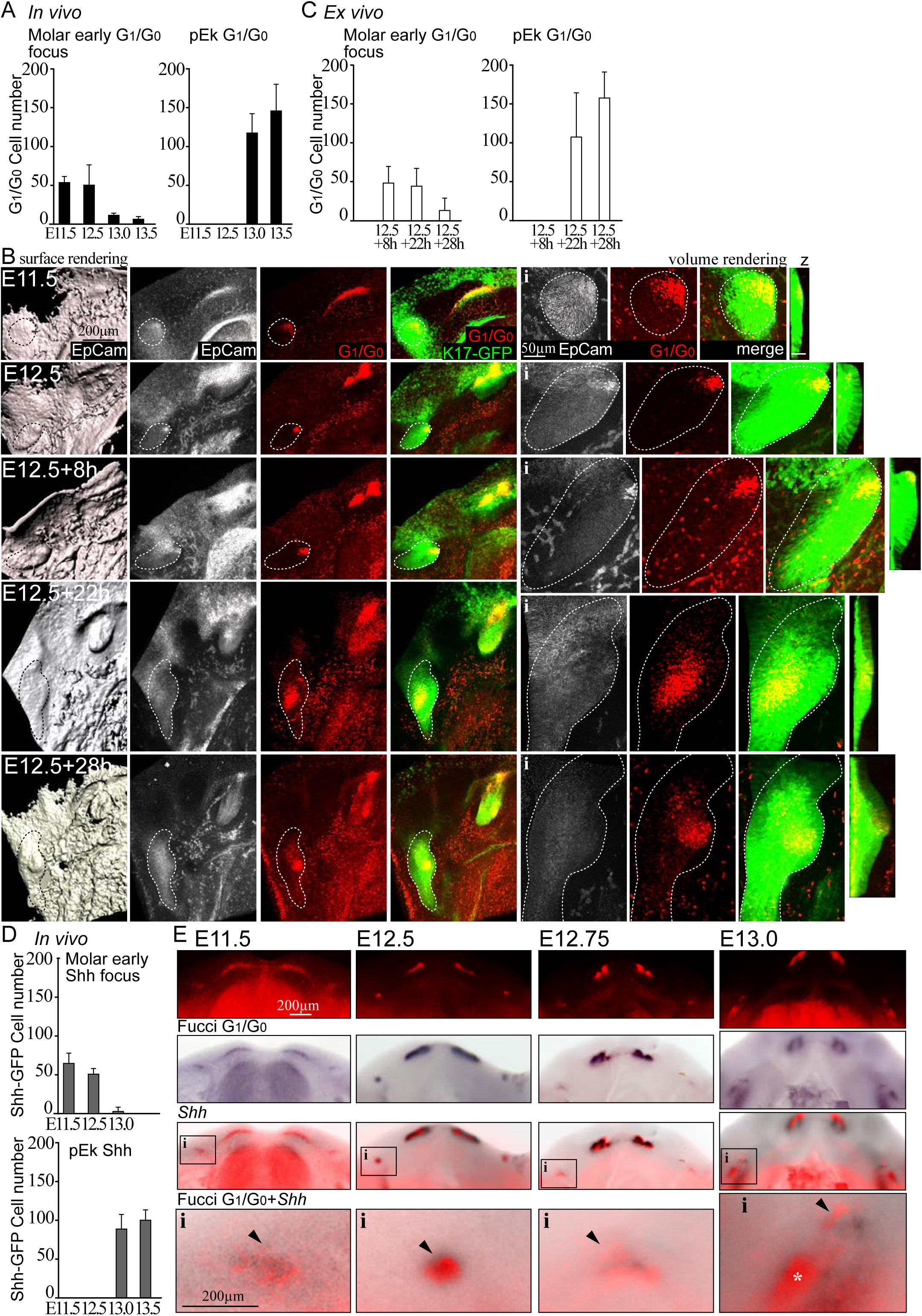
Mandible explants grown *in vitro* show same cell behavioral and morphogenetic developmental patterns as in *in vivo*. For functional live tissue imaging analyses we first verified that cultured explants developed, in respect to G_1_/G_0_ cell population dynamics and bud growth, comparably to *in vivo* development. (A) Quantification of G_1_/G_0_ cells in the developing molar placode and bud *in vivo* (embryos N_E11.5_=7, N_E12.5_=11, N_E13.0_=4 N_E13.5_=5, error bars±SEM). (B) Confocal fluorescence images of mouse embryonic mandible explants at E12.5 and grown *ex vivo* in a Trowell culture setup for 8h, 22h or 28h. Fucci fluorescent cell cycle indicator G_1_/G_0_ (Fucci G1)(red), K17-GFP epithelial placode/bud marker and immunofluorescence staining for a pan-epithelial marker EpCam (grey), epithelium (dotted line). Explants show similar morphogenesis to *in vivo* with a slight lag in timing. (C) Quantification of G_1_/G_0 cells_ in the developing molar placode and bud *in vitro* in cultured explants (embryos N_E12.5+8h_=2, N_E12.5+24h_=8, N_E12.5+27h_=7, mean±SEM) (D) Quantification of Shh-GFP cells in the IK and emerging pEK *in vivo* (embryos N_E11.5_=4, N_E12.5_=11, N_E13.0_=3 N_E13.5_=5, error bars±SEM). (E) Fucci G_1_/G_0_ fluorescence images overlaid with whole-mount DIG *in situ* hybridization with a probe specific for *Shh*. Molar IK G_1_/G_0_ focus (arrowhead), emerging pEK (asterisk).

**Related to Figure 2**.

**Figure S2.**
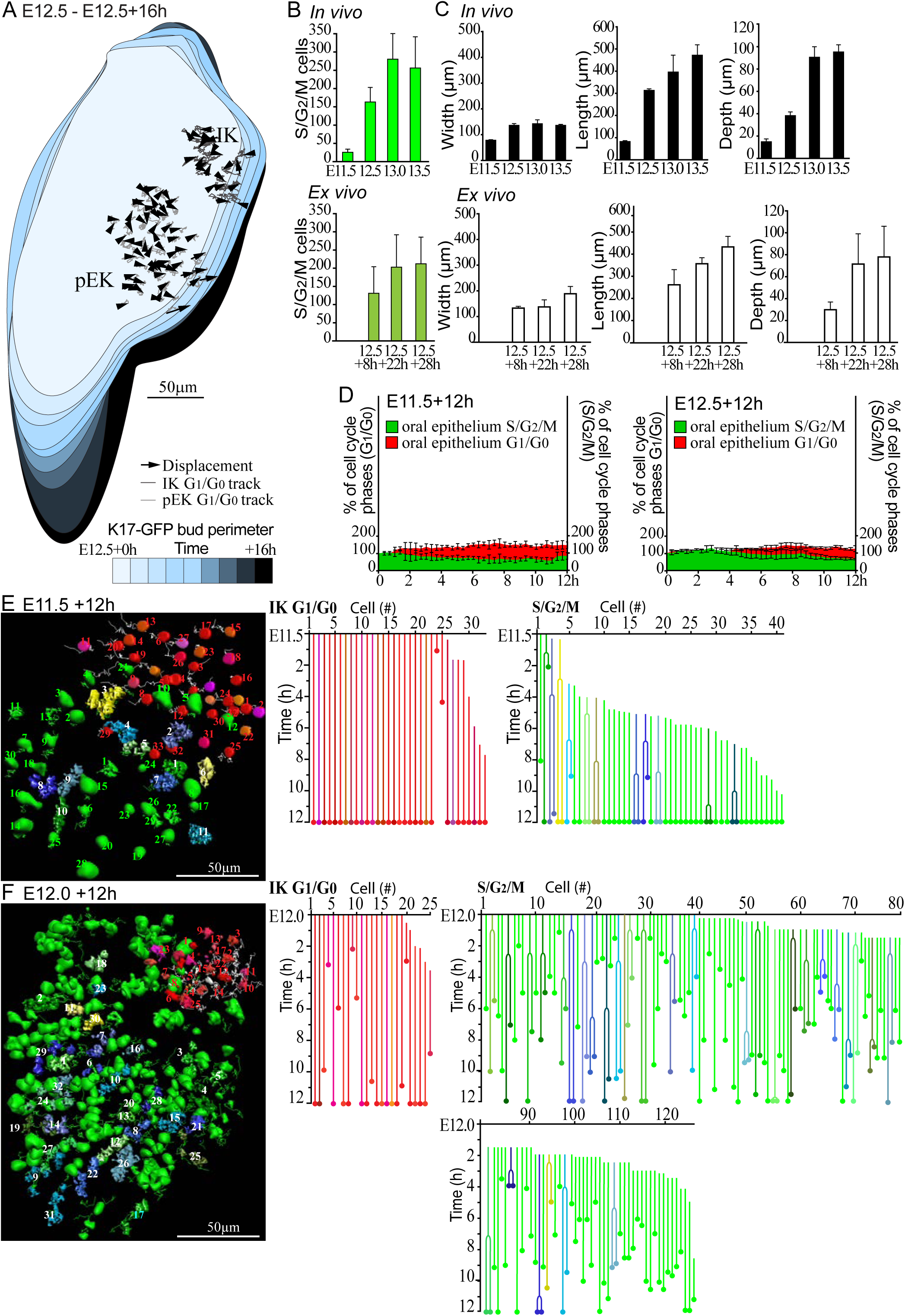
Tooth epithelial cell populations contribute differentially to the growing bud. (A) IK cells remain an integral part of the developing early molar bud. Graph shows the distribution of individual IK G_1_/G_0_ cells (cell tracks in dark grey) and emerging pEK cells (cell tracks in light grey) to the growing bud imaged from E12.5 up to 16h (image z-stacks taken at 20min intervals). Color scale represents borders of the growing tooth bud visualized with K17-GFP. (B) Quantification of S/G_2_/M cell number in developing molars *in vivo* (N_E11.5_=7, N_E12.5_=8, N_E13.0_=4, N_E13.5_=5, mean±SD) was similar to explants grown *ex vivo (*N_E12.5+8h_=4, N_E12.5+22h_=7, N_E12.5+28h_=4, mean±SD). (C) Quantification of molar bud dimensions *in vivo* (black bars) and *ex vivo* cultured explants (white bars) Dimensions in width and length were similar *in vivo* and cultured specimens with somewhat flatter bud after culturing (N_E11.5_=5, N_E12.5_=11, N_E13.0_=4, N_E13.5_=5, N_E12.5+8h_=4, N_E12.5+22h_=10, N_E12.5+28h_=7, mean±SD). (D) Quantification of cell cycle phases in oral epithelial cells of Fucci G_1_/G_0_ and S/G2/M whole mount live imaging samples from E11.5+12h (N=5, mean±SEM) and E12.5+12h (N=3, mean±SEM). (E) Surface rendering composite still image of a live tissue whole mount time lapse E11.5+12h. Tracing the contribution of individual G_1_/G_0_ and S/G_2_/M cells originating from various positions in the bud, G_1_/G_0_ (red), S/G_2_/M (green) and cell divisions (white numbers). Divisions (where cytokinesis was observed) in surface rendering shown mother and daughter cells respectively color coded. Line graph shows the contribution of individual G_1_/G_0_ and S/G_2_/M cells to the molar from E11.5+12h. (F) Surface rendering composite still image of a live tissue whole mount time lapse E12.0+12h and line graph showing the contribution of individual G_1_/G_0_ and S/G_2_/M cells.

**Related to Figure 2**.

**Figure S3.**
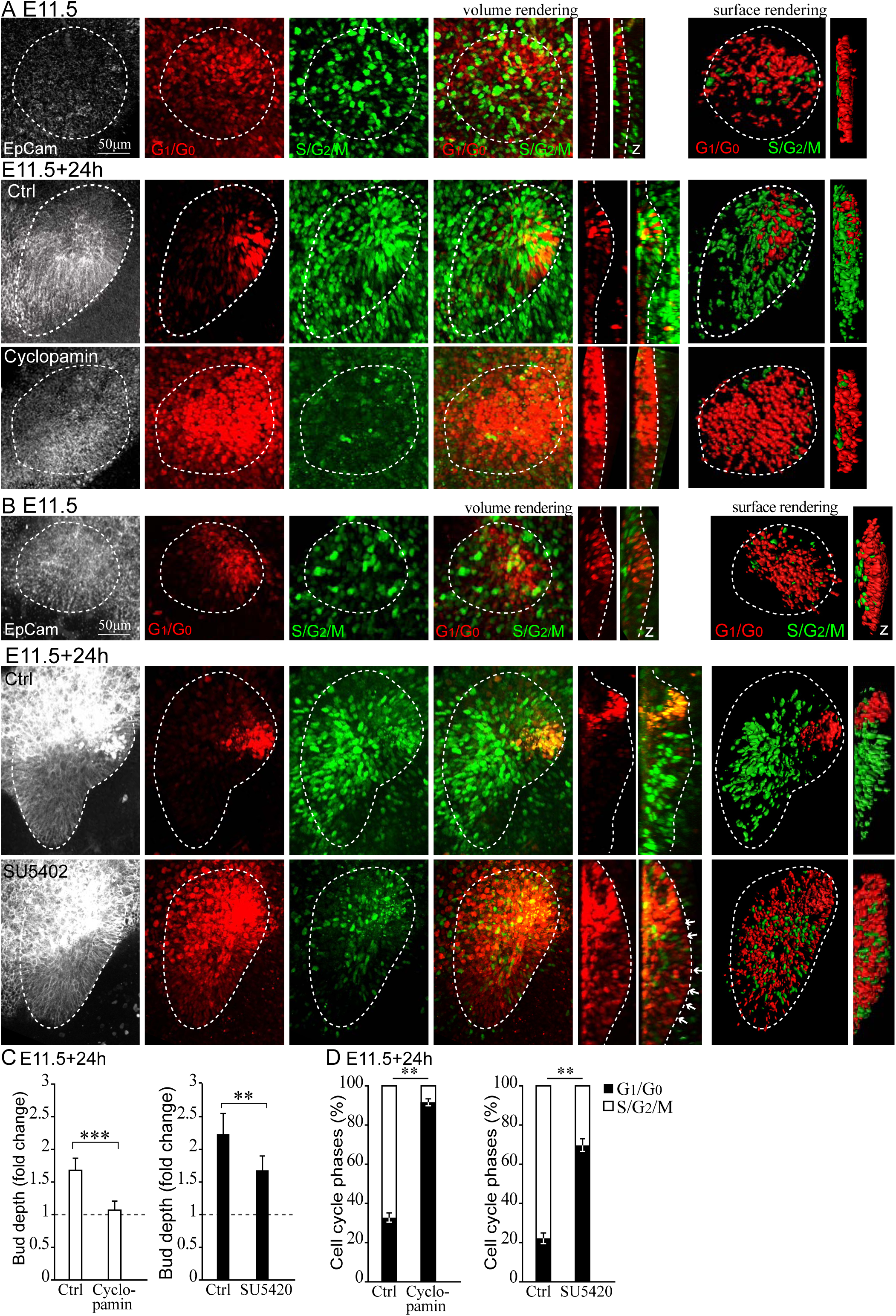
Shh is involved in bud proliferation and inhibition of Hh or Fgf signaling suppresses invagination. (A) Confocal fluorescence images of Fucci G_1_/G_0_, S/G_2_/M whole mount explants cultured E11.5+24h. Samples were treated with cyclopamine for inhibition of Shh signaling, or vehicle (Ctrl). Fucci G_1_/G_0_ (red) S/G_2_/M (green) and epithelial EpCam immunofluorescence staining (white). At E11.5, at the start of treatment the IK was emerging. After 24h in culture the control showed invagination with S/G_2_/M cells in the bud adjacent to the IK distally. Cyclopamine treatment resulted in an expansion of G_1_/G_0_ phase cells and reduced proliferation adjacent to the IK. (B) Fucci whole mount explants cultured E11.5+24h. Samples were treated with an FGFR inhibitor (SU5420) or vehicle (Ctrl). After 24h of treatment with SU5402, some proliferative cells were present but there was an increase in G_1_/G_0_ phase cells particularly in the basal population (arrows). (C) Quantification of bud depth (fold change over start of the treatment) in E11.5+24h samples treated with cyclopamine or SU5402 (N_cyclo_=6, N_SU5402_=4, mean±SEM, Mann Whitney U, *p*≤0.01**, *p*≤0.001***). (D) Quantification of G_1_/G_0_ and S/G_2_/M cells in the bud (N in all groups=5, mean±SEM, Mann Whitney U, *p*≤0.01**).

**Related to Figure 3**.

**Figure S4.**
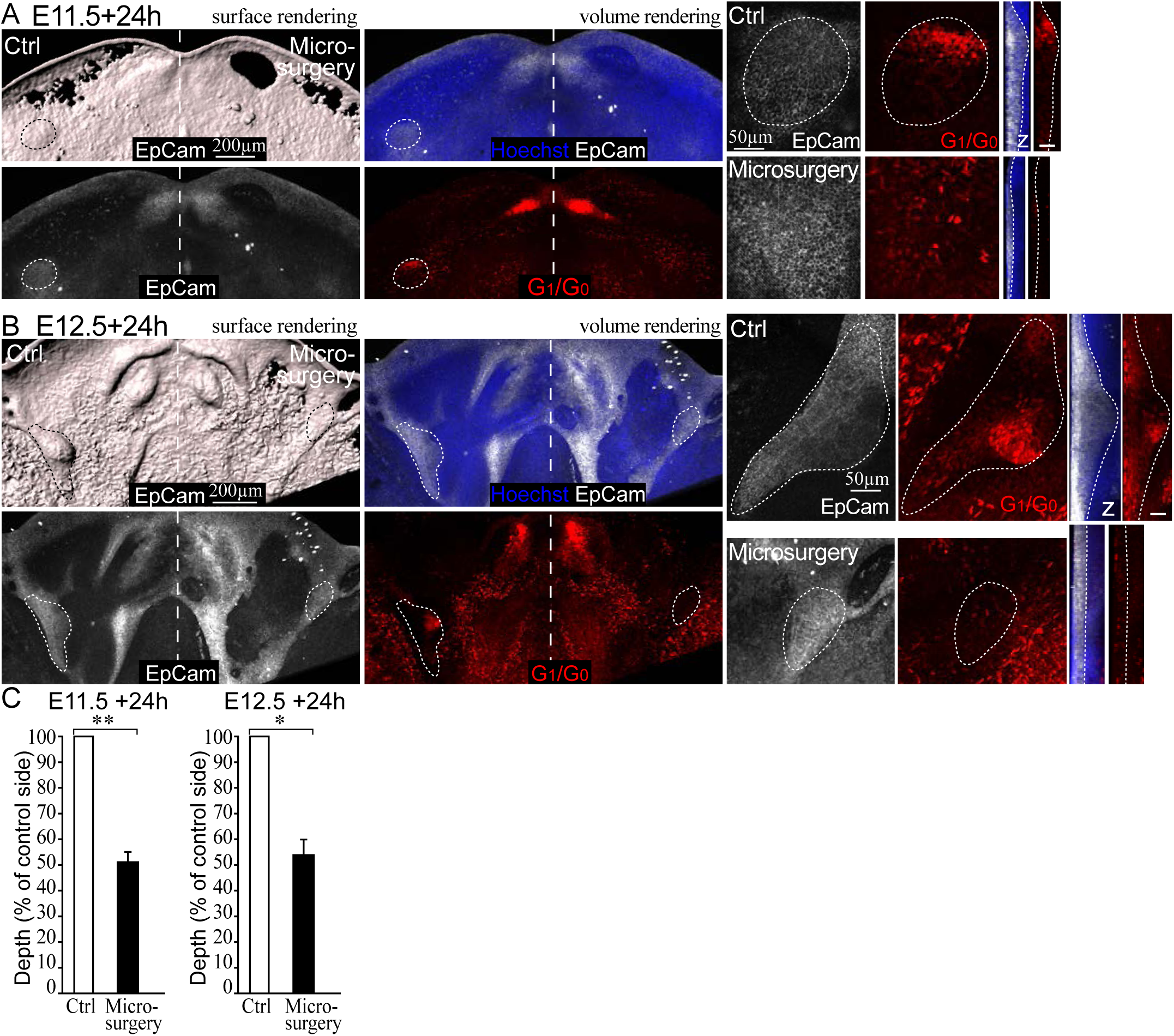
Microsurgical removal of the IK abrogates molar bud growth. Confocal fluorescence images of Fucci G_1_/G_0_ (red) whole mount explant cultures, EpCam immunofluorescence staining (white, epithelium marked with a dotted line), nuclei Hoechst (blue). (A) The molar epithelial placode was microsurgically removed at E11.5 and the tissue cultured for 24h. No G_1_/G_0_ condensate was present in the diastema and the epithelium remained flat while the control bud invaginated normally. (B) Microsurgical removal of the IK at E12.5 similarly arrested molar growth. The control side development proceeded normally to bud stage with the emerging pEK present. (C) Quantification of epithelial bud depth on the control side compared to epithelium depth in the respective area on the side where the IK was microsurgically removed in E11.5+24h and E12.5+24h explants (N_E11.5+24h_=8, N_E12.5+24h_=7, mean±SEM, Mann Whitney U, *p*≤0.05*, *p*≤0.01**)

**Related to Figure 6**.

**Figure S5.**
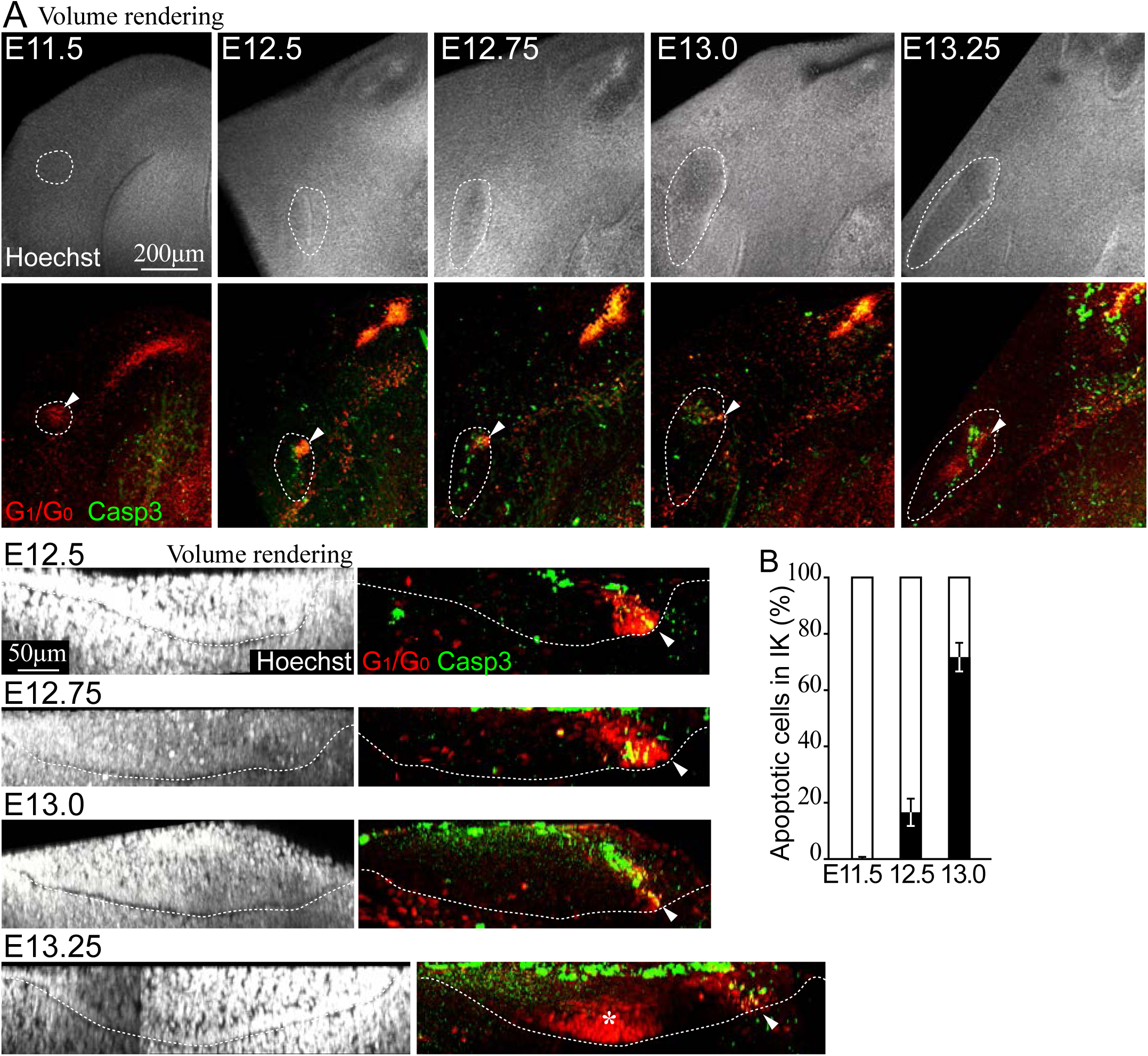
Apoptosis as a silencing mechanism of the molar IK signaling center. (A) Whole mount confocal fluorescence images of Fucci G_1_/G_0_ (red), cleaved caspase 3 immunofluorescence staining for apoptotic cells (Casp3, green). Epithelial bud (dotted line), IK (arrowhead) and presumptive pEK (asterisk), volume rendering from side view of molar. (B) Quantification of Casp3+ nuclei in the molar IK (N=12, mean±SEM).

**Related to Figure 7**.

**Figure S6.**
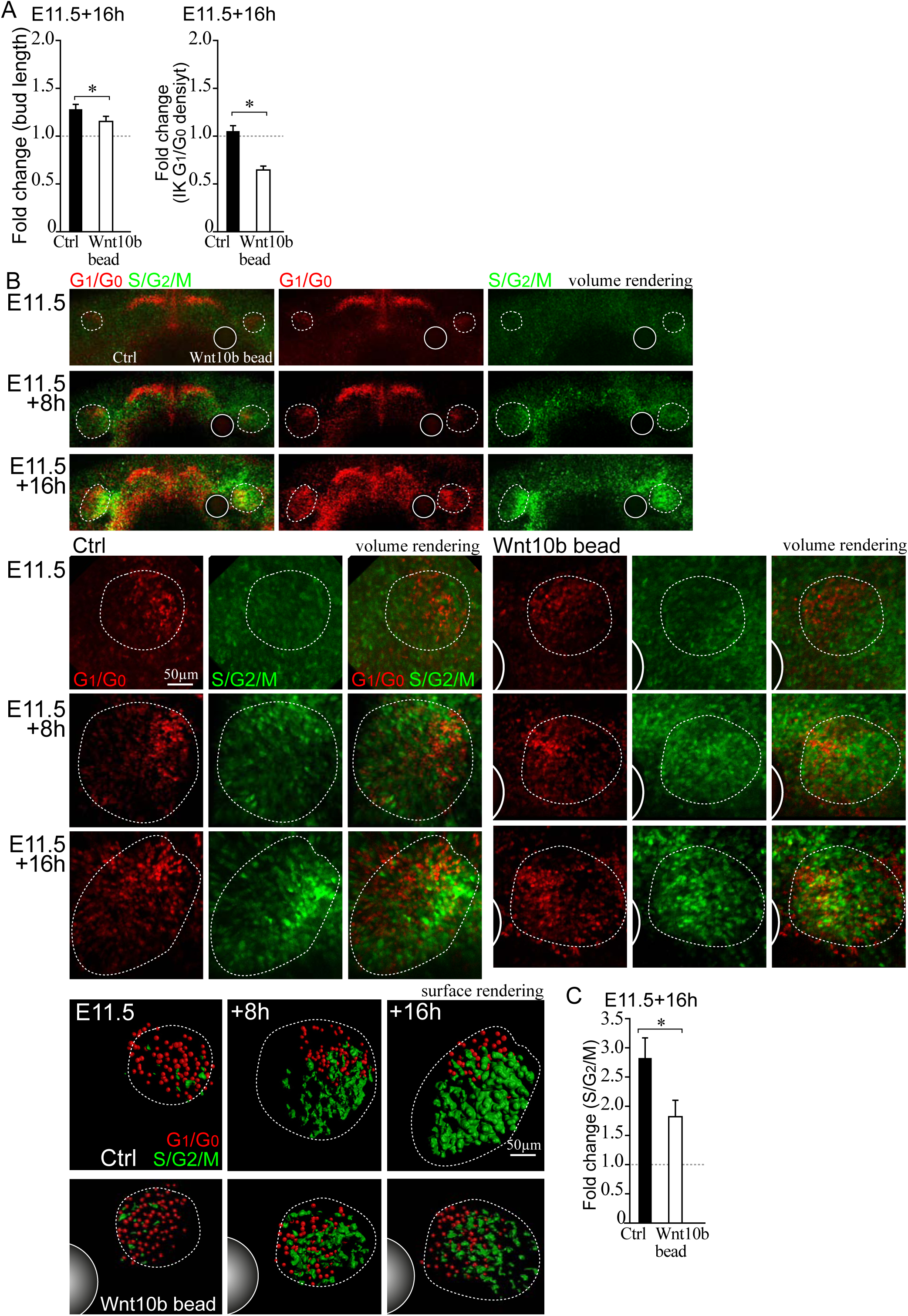
Apoptosis as a silencing mechanism of the molar IK signaling center. (A) Quantification of bud length and IK G_1_/G_0_ cell density (fold change over start of the treatment) in E11.5+16h samples with a recombinant Wnt10b protein releasing or control bead close to the IK distally on the lingual side of the placode compared to control (N=10, mean±SEM, Mann Whitney U, *p*≤0.05*). (B) Confocal fluorescence images of Fucci G_1_/G_0_ (red), S/G_2_/M (green) whole mount explant cultures, EpCam immunofluorescence staining (white, epithelium marked with a dotted line), bead (closed circle). (C) Quantification of S/G_2_/M cells at E11.5+16h (fold change over start of the treatment) in Wnt10b bead treated molar buds compared to untreated control buds (N=6, mean±SEM, Mann Whitney U, *p*≤0.05*).

## Supplemental Movie Legends

**Related to Figure 2**.

**Movie S1. Tracking individual cell G1 and S/G2/M fates in a Fucci cell cycle reporter E11**.**5+12h molar placode by live tissue confocal imaging shows placode IK cells stay in G1 phase**

Fluorescence confocal microscopy time-lapse of an embryonic mouse E11.5 whole-mount mandible explant imaged for 12h. Image stacks were taken at 20min intervals, and the playback speed here is five frames per second. The tracking of individual cell fates from the placode stage is shown as a surface rendering of the Fucci cell cycle indicator: the contribution of individual G_1_/G_0_ phase cells in the placode and initiation knot are shown color coded in red hues and S/G_2_/M cells in green. Individual cell divisions of mother cells and their daughters are color coded in blue, green and yellow shades, respectively. IK G_1_/G_0_ cells in the placode did not re-enter the cell cycle. In contrast neighboring cells, distally from the IK, frequently entered S/G_2_/M phase and cell divisions contributed to invagination locally.

**Related to Figure 2**.

**Movie S2. The IK remains an integral part of the growing tooth bud as shown in Fucci cell cycle reporter E12**.**0+12 h molars by confocal live tissue imaging**

Fluorescence confocal microscopy time-lapse of a Fucci cell cycle indicator E12.0 whole-mount mandible explant imaged for 12h. Image stacks were taken at 20min intervals, and the playback speed here is five frames per second. The tracking of individual cell fates during the early bud invagination, is shown as a surface rendering: G_1_/G_0_ phase cells are shown in red hues and S/G2/M cells in green. Individual cell divisions of mother cells and their daughters are color coded in blue, green and yellow shades, respectively. IK cells remained in G_1_/G_0_ phase and remained an integral part of the growing tooth. S/G_2_/M phase cells and cell divisions in both basal and suprabasal populations contributed to invagination and growth throughout the bud.

**Related to Figure 4**.

**Movie S3. Live imaging of Fucci cell cycle reporter shows that IK cells rearrange dynamically in the E11**.**5+12h molar placode**

Fluorescence confocal microscopy time-lapse of an embryonic mouse E11.5 whole-mount mandible explant imaged for 12h, showing the contribution of cell cycle stages to molar placode and initial budding morphogenesis on a high single cell resolution. Image stacks were taken at 20min intervals, and the playback speed here is five frames per second. The movie shows a volume rendering of cell cycle indicator Fucci G_1_/G_0_ nuclei (red) and S/G_2_/M (green). First, an overview of the mandible at the start of imaging seen from the mesenchymal side toward the epithelium, then a close up of the developing molar placode (IK, open circle) followed by both channels merged and separately. Individual cell tracks are shown as a dragon tail rendering showing a subset of twenty subsequent points in each track, IK G_1_/G_0_ (red), S/G_2_/M (green). The IK G_1_/G_0_ cells moved toward the mesial front area of the bud. The IK cells stayed in G_1_/G_0_ phase and drove proliferation locally in the adjacent cells, posterior to the knot, to initiate the invagination of the epithelium. The bud S/G_2_/M cells showed little movement.

**Related to Figures 3, 4 and 6**.

**Movie S4. Live imaging shows TCF/Lef:H2B-GFP and Fucci G1 marker expressing cells are closely juxtaposed and exhibit differential movement patterns in E11**.**5+12h molar placodes**

Fluorescence confocal microscopy time-lapse of E11.5+12h Fucci G_1_/G_0_ and TCF/Lef:H2B-GFP reporters. Image stacks were taken at 20min intervals, and the playback speed here is five frames per second. The movie shows a volume rendering of Fucci G_1_/G_0_ nuclei (red) and TCF/Lef:H2B-GFP canonical Wnt signaling reporter (green). First an overview of the mandible at the start of imaging seen from the mesenchymal side toward the epithelium, then a close up of the developing molar placode (IK, open circle) followed by both channels merged and separately. Individual cell tracks are shown as a dragon tail rendering showing a subset of ten subsequent points in each track, IK G_1_/G_0_ (red), TCF/Lef:H2B-GFP (green), both reporters (yellow). The molar placode IK G_1_/G_0_ cells specifically localized to the peripheral border formed by dental lamina cells with high TCF/Lef:H2B-GFP reporter expression (Wnt^Hi^ cells). Increasing numbers of G_1_/G_0_ cells were recruited to the IK with directional movement, toward the dental lamina Wnt^Hi^ cells. Dental lamina Wnt^Hi^ cells remained mostly localized.

**Related to Figures 3 and 4**

**Movie S5. Live imaging of E12**.**5+12h Fucci reporter shows that IK G1 cells do not contribute clonally to the primary EK in the molar**

Fluorescence confocal microscopy time-lapse of E12.5+12h Fucci explant, showing the contribution of cell cycle stages at bud stage and initiation of the pEK. Image stacks were taken at 20min intervals, and the playback speed here is five frames per second. The movie shows volume rendering of Fucci G_1_/G_0_ nuclei (red) and S/G_2_/M (green) overview of the mandible at the start of imaging (initial molar bud outlined white). In the close up of the molar IK is marked with an open circle and the location where the pEK will be initiated (marked with a closed circle). Time lapse shows volume rendering of both channels merged and separately. In track view the perimeter of the mature bud is outlined in white. Individual cell tracks are shown as a dragon tail rendering showing a subset of ten subsequent points in each track, IK G_1_/G_0_ (red), pEK G_1_/G_0_ (magenta), S/G_2_/M (green). A wave of cell proliferation contributed to rapid bud growth while the IK cells stayed in G_1_/G_0_ in the mesial part of the bud. The pEK G_1_/G_0_ cells were initiated *de novo* deep in the tip of the invaginating bud with no clonal contribution from the IK.

**Related to Figures 4**.

**Movie S6. Live imaging of Fucci and TCF/Lef:H2B-GFP reporters shows independent IK and pEK signaling centers regulating morphogenesis in E12**.**5+12h molar**

Fluorescence confocal microscopy time-lapse of Fucci G_1_/G_0_ and TCF/Lef:H2B-GFP E12.5+12h whole mount explant. The movie shows volume rendering of Fucci G_1_/G_0_ nuclei (red) and TCF/Lef:H2B-GFP canonical Wnt signaling reporter (green) visualizing the IK and emerging pEK signaling centers. Image stacks were taken at 20 min intervals, and the playback speed here is five frames per second. An overview of a volume rendering of the mandible at the start of imaging shows both channels merged and the molar bud is outlined in white. In close up view of the molar IK position is marked with an open circle and the location where the pEK is later initiated is marked with a closed circle. Time lapse shows volume rendering of both channels merged and separately. In track view the perimeter of the mature bud is outlined in white. Individual cell tracks are shown as a dragon tail rendering showing a subset of ten subsequent points in each track, IK G_1_/G_0_ (red), pEK G_1_/G_0_ (magenta), high intensity TCF/Lef:H2B-GFP (Wnt^Hi^, green), double positive (Wnt^Hi^+ Fucci G_1_/G_0_, yellow). Initially at E12.5, Wnt^Hi^ cells surrounded the G1 cells in the IK. TCF/Lef:H2B-GFP positive cells appeared throughout the bud with increase in signal intensity in the prospective pEK region followed by emergence of first G1 cells. Tracking of individual G_1_/G_0_, Wnt^Hi^, and double positive cells showed that none of these cell populations from the IK contributed clonally to the pEK; the Wnt^Hi^ and G_1_/G_0_ cells in the pEK region appeared *de novo*.

